# Experience dependence of alpha rhythms and neural dynamics in mouse visual cortex

**DOI:** 10.1101/2022.11.11.516158

**Authors:** Pouria Riyahi, Marnie A. Phillips, Nathaniel Boley, Matthew T. Colonnese

**Affiliations:** The George Washington University, Department of Pharmacology and Physiology, Washington, D.C. 20052; The George Washington University School of Medicine, Department of Biomedical Engineering, Washington, D.C. 20052; The George Washington University School of Medicine, Institute for Biomedical Sciences, Washington, D.C. 20052

## Abstract

The role of experience in the development and maintenance of emergent network properties such as cortical oscillations and states is poorly understood. To define how early-life experience affects cortical dynamics in adult visual cortex, we examined the effects of two forms of blindness, bilateral loss of retinal input (enucleation) and degradation of visual input (eyelid-suture), on spontaneous activity in awake head-fixed mice of both sexes. Neither form of deprivation fundamentally altered the state-dependent regulation of firing-rates or local field potentials. However, each form of deprivation did cause a unique set of changes in network behavior. Laminar analysis revealed two different generative mechanisms for low-frequency synchronization, one prevalent during movement, the other during quiet-wakefulness. The former was absent in enucleated mice, suggesting a mouse homolog of human alpha oscillations. In addition, neurons in enucleated animals were less correlated and fired more regularly, but showed no change in mean firing-rate. Chronic lid-suture decreased firing rates during quiet-wakefulness, but not during movement, with no effect on neural correlations or regularity. Sutured animals showed a broadband increase in dEEG power and an increased occurrence, but reduced central frequency, of narrowband gamma oscillations. The complementary--rather than additive--effects of lid-suture and enucleation suggest that the development of these emergent network properties does not require vision but is plastic to modified input. Our results suggest a complex interaction of internal set-points and experience determines the expression of mature cortical activity, with low-frequency synchronization being particularly susceptible to early deprivation.

**SIGNIFICANCE STATEMENT:** The developmental rules that guide how cortex balances internal homeostatic set points with external inputs to establish the emergent network level dynamics critical to its function are unclear. Using multiple methods of early deprivation, we show that the development of dynamics in mouse visual cortex is not dependent on the type of input. Rather, specific neural rhythms, firing-rate set points, and neural correlations are differentially modified by experience. Our deprivations identify one specific rhythm as a likely homolog to human alpha and suggest a mechanism for its loss in blindness. Our results advance our understanding of the regulatory mechanism leading to normal cortical processing, which is altered in blindness and multiple neural disorders.

## INTRODUCTION

Computations in cortical circuits depend not only on the activity and organization of the inputs, but also on the interaction of inputs with ongoing and dynamic ‘endogenous’ activity generated in the thalamocortical circuit (Harris and Thiele, 2011; Poulet and Crochet, 2019; McCormick et al., 2014). This ongoing activity is reflected in emergent network behaviors such as oscillations of different frequency bands that reflect the underlying patterns of synchronization and are strongly modulated by behavioral state (Uhlhaas et al., 2008; Buzsáki and Draguhn, 2004). The rules by which ongoing spontaneous network dynamics and their regulation by state are established and maintained during development are poorly understood. In the visual system, deprivation profoundly alters the organization of inputs and local circuits (Hooks and Chen, 2020; Wang et al., 2021) leading to persistent visual deficits more severe than would be predicted by the response properties of neurons alone (Mitchell and Maurer, 2022; Kiorpes, 2016). Visual deprivation results in substantial remodeling of the visual cortex resulting in increased cross-modal input, increased blood flow and energy usage, and greater responses to non-visual stimuli (Merabet and Pascual-Leone, 2010; Röder and Kekunnaya, 2022; Burton, 2003; Veraart et al., 1990; Mishina et al., 2003). The cortex homeostatically regulates neuronal excitability and network properties (Wen and Turrigiano, 2024), making it resilient to large disruptions in input. However, the limits of homostatic mechanisms, the developmental plasticity of the set points, and the interaction between sensory deprivation, homeostasis of firing rates, network interactions, and cortical oscillations are poorly understood. In humans congenital blindness has a mixed effect on cortical oscillations, profoundly reducing alpha activity in visual cortex (Adrian and Matthew, 1934; Jeavons, 1964; Noebels et al., 1978; Lubinus et al., 2021), while leaving other bands relatively intact (Hawellek et al., 2013). Visual deprivation affects slow-wave activity during sleep in mice (Miyamoto et al., 2003), but cortical activity changes similar to those observed in humans during wakefulness have not been described.

Head-fixed mice on a treadmill display changes in cortical oscillations similar to those of primates during attention shifts (Harris and Thiele, 2011). These include suppression of large amplitude slow-waves and increased gamma-band power, increased firing-rates, and visual-response gain with increasing arousal (McGinley et al., 2015). The initial emergence of adult-like network dynamics in thalamus and cortex occurs just before eye-opening (Colonnese, 2014; Murata and Colonnese, 2018) and is unaffected by enucleation (Riyahi et al., 2021). Neuronal firing-rates and gamma oscillations mature in the 3rd and 4th postnatal weeks (Hoy and Niell, 2015; Shen and Colonnese, 2016)) and this is delayed by dark-rearing (Chen et al., 2015).

Here we examined the effect of two forms of early-life blindness on the establishment and maintenance of cortical dynamics. Bilateral enucleation at P6 or P13 allows us to compare the effect of retinal input removal before and after the onset of mature dynamics (Riyahi et al., 2021), without the significant thalamic and cortical rewiring caused by earlier loss of retinal inputs in mice (Olavarria and Hiroi, 2003; Golding et al., 2014). Binocular eyelid suture initiated before eye-opening deprives the animal of pattern vision and the ability to guide behavior visually, while preserving luminance detection and circadian rhythms (Kampf-Lassin et al., 2011), similar to bilateral cataracts. We find a remarkable resilience of visual circuit dynamics to lifelong visual deprivation. We also identify several network behaviors modified by reduced visual experience, including a loss of low-frequency activity that suggests the mouse homolog of human alpha rhythms.

## Methods and Materials

### Animal care

Animal care and procedures were in accordance with *The Guide for the Care and Use of Laboratory Animals*(NIH) and approved by the Institutional Animal Care and Use Committee at The George Washington University. Postnatal day(P)0 is the day of birth. C57BL/6 were obtained from Hilltop Lab Animals(Scottsdale, PA) as timed pregnant females, and kept in a designated, temperature and humidity-controlled room on a 12/12 light/dark cycle and examined once per day for pups. Large litters were reduced to 6-8 pups. Each litter was split approximately evenly into experimental or control animals, pups were not sexed, and all pups within the litter were used.

### Surgical Procedures

For bilateral enucleations, carprofen (5 mg/kg) in saline was injected 1 hour before surgery to reduce pain and inflammation. Surgical anesthesia was induced with 3% isoflurane vaporized in 100% O_2_, verified by toe-pinch and respiratory rate and effort. Thermal support was provided during surgery. An incision was made in the eyelid (P6) and the globe of the eye was removed using forceps. For older animals (P13), the eyelid attachment had thinned enough to enable manual opening of the eyelid with only light pressure. The eye socket was filled with sterile surgical foam (GelFoam) and the eyelid closed using a tissue adhesive (Vetbond). Pups were post-operatively monitored and received follow-up injections of carprofen daily for 2 days. Sham control littermates received identical treatment, including eyelid puncture with the tip of a suture needle, without enucleation or GelFoam. For bilateral eyelid suture, on P11 animals were anesthetized and prepared for surgery as above except that hair removal was also necessary. This was achieved by placement of Vetbond over all pups’ eyelids 2-3 days prior to surgery. Dams removed the vetbond during grooming and this also removed the underlying hair without tearing or chemical burn of the delicate eye tissue. To suture the eyes, an incision was first made with iris scissors to separate the eyelid. The cut ends of the incision were opposed manually and the incision was fixed using running horizontal mattress microsutures (Size 9-0, AROSurgical T06A09N14-13) to bring the cut ends neatly together. In this way, the eyelid suture is held in place long-term by the healed incision as dams remove the sutures within days. Pups received follow-up analgesia and sham littermates were treated similarly. The mice were observed daily for signs of incomplete fusion and any pups with any eyelid opening were removed from the study. Eyes were dissected post-mortem and examined for lens and corneal clarity and found to be clear, albeit with some scar tissue formation under the healed incision. Corneas were also examined with fluorescein (2% sodium USP) to assay cornea health and no wounding was observed.

For headpost placement, carprofen(5 mg/kg) in saline was injected 1 hour prior to surgery to reduce pain and inflammation. Surgical anesthesia was induced with 3% isoflurane vaporized in 100% O_2_, verified by toe-pinch and respiratory rate and effort, then reduced to 1.5-3% as needed by monitoring breathing and toe pinch response. An electrical heating pad(36°C) provided thermoreplacement. For attachment of the head-fixation apparatus, the scalp was excised to expose the skull, neck muscles were detached from the occipital bone, and the membranes were removed from the surface of the skull. Topical analgesic was applied to the incision(2.5% lidocaine/prilocaine mix, Hi-Tech Pharmacy Co., Inc., Amityville NY). The head-fixation apparatus was attached to the skull with grip cement(Dentsply, Milford DE) over Vetbond™ tissue adhesive(3M). The fixation bar consisted of a custom-manufactured rectangular aluminum plate with a central hole for access to the skull. After placement, the animal was maintained with 0.5-1% isoflurane until the dental cement cured, after which it recovered on a warming table. Pups were post-operatively monitored and received follow-up injections of carprofen daily for 2 days. On the 3rd-5th day following surgery, they began habituation to the recording apparatus. Habituation consisted of two days of exposure to the recording apparatus. Animals were placed in fixation under isoflurane anesthesia and given 10(day 1) and 30(day 2) minute exposures after recovery from anesthesia, including visual stimulation as required.

### In vivo electrophysiology

Mice were recorded when they were 60-100 days old. After two days of habituation the animals were placed in the setup as before, but a craniotomy and electrode placement was performed while under anesthesia. For the craniotomy, the skull was thinned using a surgical drill, and small bone flaps were resected to produce a ∼150-300 µm diameter opening. Monocular visual cortex was targeted by targeting the region with the least vascular density 2.0-2.5mm lateral and 0.0-0.5 mm rostral to lambda. All recordings were made using a single shank, 32 channel array arranged in two parallel lines of contacts(A1×32-Poly2-5mm-50s-177, NeuroNexus Technologies, Ann Arbor MI). The electrode penetrated the brain orthogonally to the surface and was advanced to a depth of 500-800µm using a stereotaxic micromanipulator until the top channels touched the cerebral-spinal fluid. Isoflurane was withdrawn and the animal allowed to recover for 60 min before recordings. In the absence of visual stimulation, the room was completely dark(<0.01 Lumens).

For the enucleation experiments animals were recorded for 30 min in the dark. For the eyelid suture experiments, animals received visual stimulation from an LED monitor centered 30 cm from the contralateral eye subtending 120° of the contralateral visual field. Visual stimuli were displayed using custom software in Matlab using PsychToolbox (Brainard, 1997; Pelli, 1997). Stimuli consisted of 3 alternating blocks, each 60 secs, repeated for 30 min (10 exposures to each block). The blocks were “Black” consisting of total darkness, “Gray” consisting of a gray screen 130 Lumen/m^2^ at the eye, and “Noise” with the same total luminance divided into blocks, with ∼10 degrees of visual space randomly assigned to 0, 0.25, 0.5, 0.75 and 1 of the screen maximal luminance. The grids were updated at 5Hz. While total luminance was constant, local light intensity varied by +/−10 Lumens/m^2^ with each grid change.

### Data acquisition and analysis

Movement was measured by optical readout of displacement of a circular treadmill, digitized at 1KHz and recorded with a custom Matlab macro. For the neural signal, data was amplified 192V/V and digitized at 30 kHz on a SmartLink Headstage and recorded with the SmartBox 1.0(Neuronexus, Ann Arbor MI). Recordings were imported in Matlab using Neuronexus supplied macros and custom code. Depth (d)EEG/LFP signals were derived by down-sampling wide-band signals to 1 kHz after application of 0.1-350Hz zero-phase low-pass filter. For single-unit analysis, channels with high noise (typically those outside the brain or bad contacts) were manually eliminated and the remaining channels saved as binary structures for spike sorting using Kilosort 1.0 (Pachitariu et al., 2016). Clusters were identified using the default parameters, except ops.Th = [ 3 6 6]. Clusters were manually merged and sorted into noise, multi-unit, and single-unit clusters using the GUI Phy (Rossant et al., 2016). Following sorting, each good cluster was evaluated for inclusion as Regular or Fast-spiking using a modification of (Vinck et al., 2015). Good clusters were separated by k-mean clustering using the normalized spike amplitude of the mean waveform at 8/32 and 60/32 of a ms after the spike peak. The number of clusters was set to 3 and the two clusters with longest spikes were assigned to the regular spiking group and the other to the fast-spiking group.

All analyses were performed in Matlab. Spike-times were binned at 1 ms, and aligned with dEEG signals. Animals were evaluated for inclusion based on the following criteria:(1) at least 10 well-isolated ‘good’ units;(2) at least 10% and less than 50% time moving; (3) dEEG displaying normal relationships between layers, including elevated high(20+) frequency power relative to deep layers (Senzai et al., 2019) and prominent positive peaks at the surface, (4) absence of spreading depression and paroxysmal activity, (5) absence of eyelid separation for sutured animals. Spreading depression was identified as a sawtooth oscillation that spread slowly from surface to deep layers over 10-20 seconds and was followed by 1 or more minutes of silence. Paroxysmal activity was defined by the presence of 5 more seconds of high-amplitude(>3 fold above the maximal envelope for the remainder of the recording) spikes. In total 15/48 animals were eliminated from the enucleated experiment (9/19 sham; 6/29 Enucleated); and 12/29 animals(5/12 sham; 7/17 sutured) were eliminated from the suture experiments.

Layer assignment for each contact was made by analysis of the MUA and dEEG signals using Senzai et al., 2019 as a guide. In a subset of early animals, this assignment was verified by the reconstruction of the electrode track. In each animal, the surface/L1 was assigned as the contact with a reversal in the dEEG signal and consequent reduction in spectral power. Additionally, this site was above the top contact with action potentials. Border between superficial and deep layers was assigned as the channel located at the bottom of the elevated high-frequency power (>20Hz) observed in L2-4 (Fig. 1B). This position aligned with the bottom of the initial visually evoked sink in sighted animals (Fig. 1B). A band of consistent, large-amplitude spiking was often, but not always, observed ∼100 µm below the L4 channel (Senzai et al., 2019). For assignment of the dEEG signal, the contact with the least movement artifact <200 µm above the superficial/deep division was selected. For Figure 9, L6 LFP was chosen as the contact with no artifact at least 300-400 µm below L4 and deeper than the region of low power (L5). For analysis, recordings were divided into 500ms windows, and the maximum velocity of the treadmill, firing rates for all neurons(expressed in Hz), and spectral power of the dEEG was calculated. For the dEEG signal, the channel no more than 200 µm from the bottom of L4 with the least artifact was selected. Spectral decomposition of the dEEG signal used the multi-taper method (Mitra and Bokil, 2007) using a ‘time-bandwidth product’ 3 and ‘number of tapers’ 5. Spectral power was whitened by the multiplication of power by frequency. Each 500 ms window was sorted into Moving and Stationary conditions based on the presence or absence of movement of the circular treadmill. Mean spectra for each animal were calculated for Moving and Stationary periods, as were mean firing rates for each neuron. Spiking regularity(LvR) was calculated by the method of (Shinomoto et al., 2009) for each cluster across the entire recording. Calculation of spike-rate covariance followed the method of (Renart et al., 2010). Each single-unit raster was convolved by a normalized kernel that was the sum of a rapid time-window(J) and a negative longer window(T, 4xJ). All analysis except for the effect of J (Fig. 7F), was made with J=160ms. For the isolation of 3-6Hz oscillations we identified periods of high amplitude negative potentials in the superficial dEEG in which at least 3 negative peaks greater than 3 standard deviations below the mean negative potential occurred within 1 s. These periods were further restricted to require at least 20 ms with no spiking(combined multi and single unit) between each high-amplitude spike. This method was most consistent with identifying all periods of visible 3-6Hz oscillation occurring after movement. For the isolation of individual low-frequency events (Fig. 9), the LFP signal was 2-12 Hz band-pass filtered and the negative and positive peaks for each channel were identified using the ***findpeaks*** function. Peaks in in the highest tertile with greater than 100ms separation were considered events for analysis.

**Figure 1.**
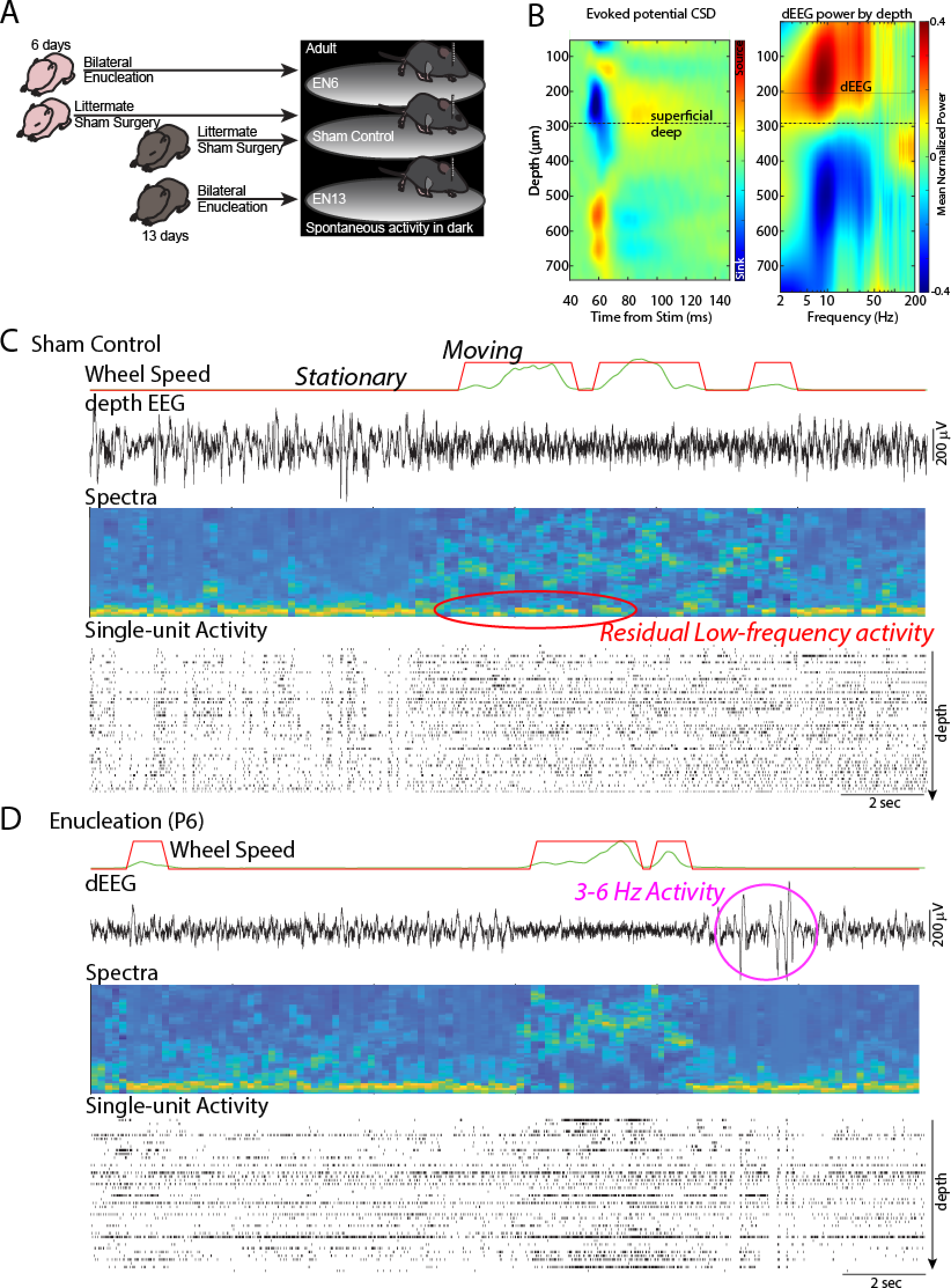
Effect of bilateral enucleation on cortical dynamics. **A.** Experimental design. Mouse pups underwent bilateral enucleation at 6 days old (EN6), before the development of continuous background activity and adult-like cortical states, or at 13 days old (EN13), before eye-opening but after the initial emergence of background cortical activity and states(Shen and Colonnese, 2016; Riyahi et al., 2021). Littermates of each group received sham surgery and were recorded on interleaved days. Activity in the monocular visual cortex was recorded in adult head-fixed, awake animals allowed free movement on a circular treadmill in the dark. **B.** Laminar identification. Left: current source density of the mean flash-evoked response in a Sham animal; Right: mean-normalized depth spectra of spontaneous activity in the dark from the same animal. Separation between superficial (L1-4) and deep (L5-6) was determined as the transition point between high and low 5-20Hz power. This corresponded to the bottom of the early evoked potential in sighted animals and the band of high-frequency (100Hz+) power in layer 5 present in most animals. The depth EEG (dEEG) for each animal was calculated from the best electrode 100-200 superficial to the dividing line. **C.** Representative recording from sham control littermate. Aligned time-courses show running disk velocity (green) and binarized threshold (red) dividing session into Stationary and Moving periods; local field potential (dEEG) from layer 4, spectral decomposition of dEEG signal on log(power) color scale; and raster plot of isolated single-unit firing arranged from surface at the top to layer 5 below. Red circle marks periods of residual low-frequency activity during moving periods which were observed to be absent in the enucleated animals. **D.** Representative recording from EN6 animal. Note qualitatively similar network dynamics including a decrease in EEG and unit synchronization and concomitant increase in firing during movement periods, while low-frequency activity is reduced during movement. Purple circle shows periods of large amplitude ‘3-6 Hz oscillations’ that occurred more frequently in the enucleated groups.

### Statistical Procedures

All data are mean ± SEM except for the cumulative distributions of spike rates which are 95% confidence intervals and the means of firing rates which are standard deviation. The statistical test applied is noted along with results. Statistical tests were applied in Matlab using the inbuilt functions (‘ttest2’, ‘anovan’). Significance statements regarding normalized frequency power distributions were calculated by permutation analysis following the permutation test method of (Cohen, 2014), using custom macros. Frequency resolution was 1 Hz from 1-100 Hz. The significance threshold was p<0.05.

## Results

### Effects of enucleation on cortical dynamics

To examine the requirement for the primary sensory input in establishing and calibrating neural network dynamics of visual cortex, we made acute, extracellular recordings of the local field potentials (dEEG) and action potentials using single shank polytrodes from the contralateral monocular region of the primary visual cortex of awake, head-fixed adult mice. The mice were divided into four groups: P6 enucleated (EN6, n=13) and their Sham surgery littermate controls(n=5), and P13 enucleated (EN13,n=10) and their Sham littermate controls(n=5)(Fig. 1A). No significant(P>0.5 for all comparisons) or visible differences were observed for any measurement between the two sham groups and they were consolidated into a single Sham control group for analysis. To examine the state-dependence of cortical dynamics, recordings were performed while the mice were allowed free movement on a circular treadmill. To equalize the visual input to the groups, all recordings were performed in complete darkness (Fig. 1A). There was no significant difference for any group in the time spent moving (Sham 6.9+/−0.9% time moving, EN6 5.6+/−0.4%, EN13 5.1+/−0.6% (SEM), ANOVA p = 0.12 for effect of group), in movement duration (mean bout duration: Sham 855+/−118ms, EN6 1053+/− 83ms, EN13 888+/−80ms, ANOVA =0.23) or in running speed (mean speed during movement bouts: Sham 252+/−25 arbitrary units, EN6 288+/−26, EN13 268 +/−14, ANOVA p = 0.56). We further examined the distribution of bout lengths and speed but observed no differences between groups. Thus the first-order organization of movement vs quiet wakefulness within our paradigm was not affected by enucleation.

To quantify the behavior of the cortical circuit we used two primary measures: (1) dEEG from superficial layers (L3/4 border), which reveals both the low and high-frequencies that define the behavioral state (Senzai et al., 2019)(Fig. 1B) and (2) action potentials sorted into single-unit clusters throughout the depth of cortex. Single-unit clusters were divided into regular and fast-spiking units by conventional means (Vinck et al., 2015).

Qualitatively, enucleation at either age did not change the patterns of cortical network activity visible on a treadmill (Fig. 1C-D). EN6 and EN13 animals both exhibited spontaneous background activity in the dark that was similar to the Sham control and to that described previously for head-fixed mice (Niell and Stryker, 2010). Specifically, periods with no movement (Stationary) were dominated by large amplitude, synchronous, slow (below 10Hz) fluctuations of the dEEG and neural firing. Periods of movement were associated with an increase in neural firing rates, a reduction of the firing-rate variance, and a decrease in the total dEEG amplitude, reflecting a drop in the large low frequencies and an increase in small high frequencies (20 + Hz). Visual observation of multiple animals suggested two differences between Enucleated and Sham control animals: 1) the low amplitude, high-frequency dEEG observed during movement was flatter and more consistent in enucleated animals, and 2) many enucleated animals had an increased incidence and size of very high amplitude 3-6 Hz oscillations that followed the cessation of movement (Fig. 1C).

To quantify the dEEG modulation by state we calculated frequency power of the LFP from a contact in the superficial layers in one-second windows, divided into Stationary and Moving periods based on the treadmill velocity(Fig. 2A). During Stationary periods the dEEG power was not significantly different between groups for any frequency. During Moving periods both EN6 and EN13 displayed significantly lower power from 4 to 22Hz (24Hz for EN6) than Sham, confirming our visual impression that the dEEG flattens more during movement in enucleated animals. To examine the effect of movement on dEEG power we calculated the proportional change for each frequency band during Moving periods with Stationary periods as baseline. This analysis showed a significantly greater decrease (or reduced increase for frequencies above 20Hz) in power for frequencies 2-26 Hz for EN6 and 2-36 Hz for EN13. In total, these data suggest that the regulation of cortical excitability by arousal and movement (Vinck et al., 2015) is not altered by enucleation occurring either before these circuits develop (EN6) or immediately after they emerge (EN13). However, in the absence of the eyes, low-frequency oscillations experience greater suppression during movement. Interestingly this reduction in low-frequency power was limited to movement periods and was not apparent during stationary periods when these rhythms are most prevalent.

**Figure 2.**
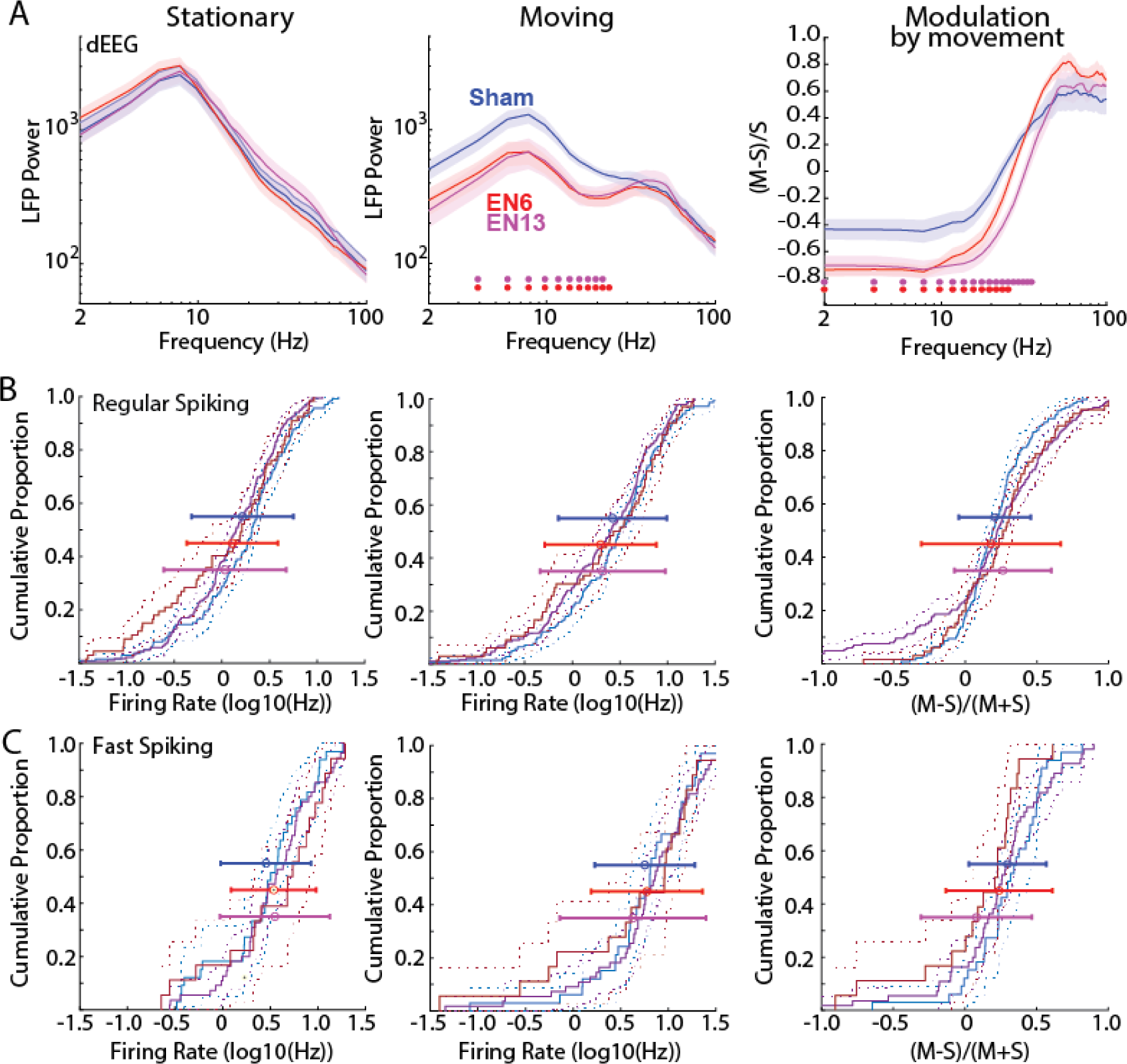
Effect of bilateral enucleation on EEG power and firing rates. **A.** Population mean and standard deviation for dEEG power for Sham (blue), P6 Enucleated(EN6; red) and P13 enucleated(EN13; purple). Power during periods of Stationary (left) and Moving (middle) periods and the proportional change during movement(Modulation by movement, right) are shown, with frequencies of significant difference(permutation test) from Sham shown by color coded dot for each experimental group. **B.** Cumulative distribution of single-unit firing-rates for regular spiking (presumptive excitatory) neurons during Stationary and Moving periods and for the modulation by movement. Solid lines show distribution and dotted lines the associated 95% confidence interval. Error bars show population mean and standard deviation for all neurons. Firing rates were not significantly different for any group (Table 1). **C.** Cumulative distribution for fast-spiking (presumptive inhibitory) neuron firing-rates.

**Table 1.**
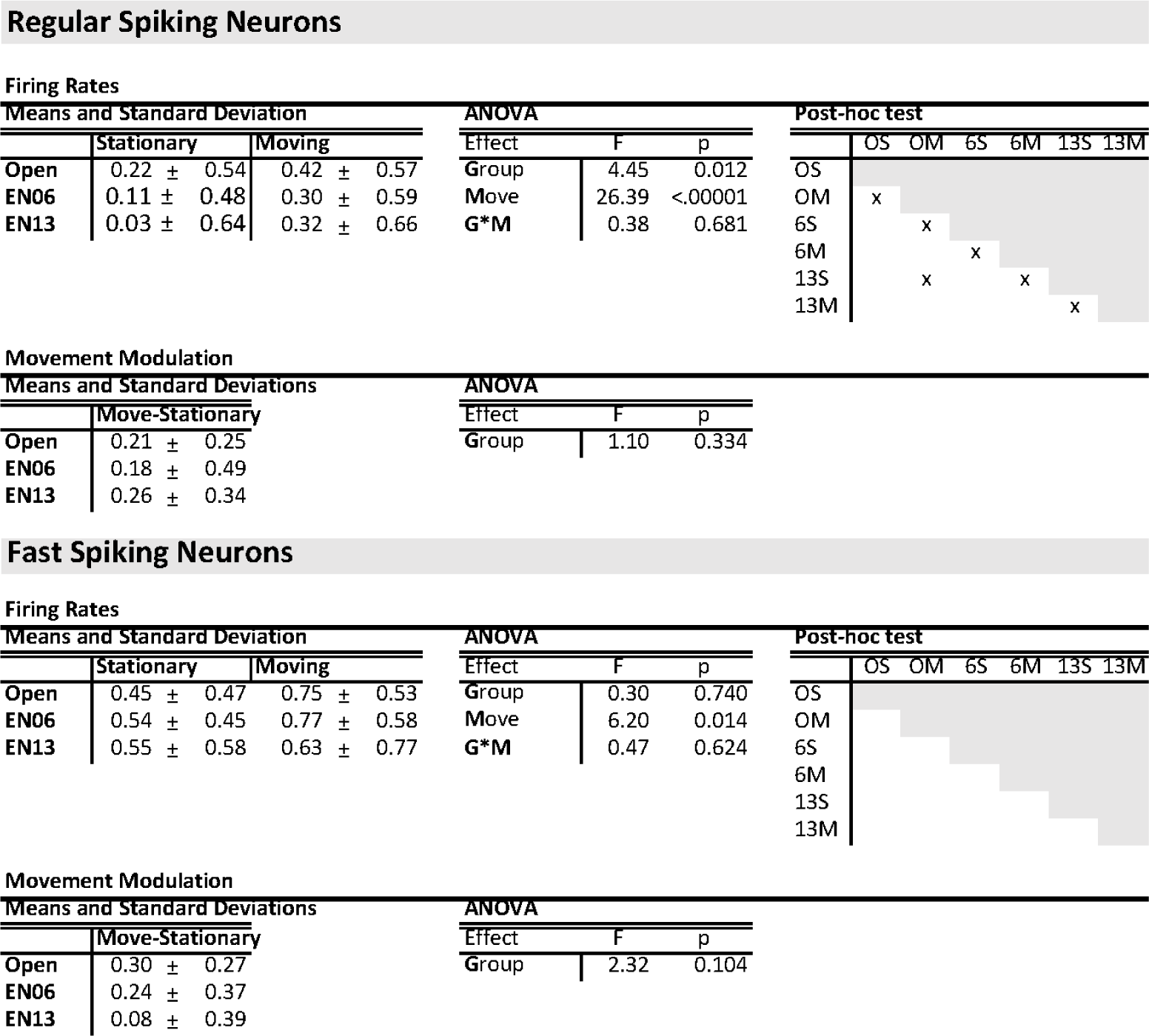
Single-Unit firing-rates following enucleation. Means and standard deviations, two-factor ANOVA and post-hoc test results for data displayed in **Fig. 2**. “Open” designates the Sham enucleated animals. For the ANOVA G*M shows the interaction between Group and Movement state. The post-hoc test results show p<0.05 by Tukey HSD, O/6/13 denote treatment group, S/M denote movement state (**S**tationary/**M**oving).

Firing Rates
| Means and Standard Deviation |  |  |  | ANOVA |  |  | Post-hoc test |  |  |  |  |  |  |  |
| --- | --- | --- | --- | --- | --- | --- | --- | --- | --- | --- | --- | --- | --- | --- |
|  | Stationary |  | Moving |  | Effect | F | p |  | OS | OM | 6S | 6M | 13S | 13M |
| Open | 0.22 ± | 0.54 | 0.42 ± | 0.57 | Group | 4.45 | 0.012 | OS |  |  |  |  |  |  |
| EN06 | 0.11 ± | 0.48 | 0.30 ± | 0.59 | Move | 26.39 | <.00001 | OM | x |  |  |  |  |  |
| EN13 | 0.03 ± | 0.64 | 0.32 ± | 0.66 | G*M | 0.38 | 0.681 | 6S |  | x |  |  |  |  |
|  |  |  |  |  |  |  |  | 6M |  |  | x |  |  |  |
|  |  |  |  |  |  |  |  | 13S |  | x |  | x |  |  |
|  |  |  |  |  |  |  |  | 13M |  |  |  |  | x |  |

Movement Modulation
| Means and Standard Deviations |  |  |  | ANOVA |  |  |
| --- | --- | --- | --- | --- | --- | --- |
|  | Move-Stationary |  |  | Effect | F | p |
| Open | 0.21 | ± 0.25 |  | Group | 1.10 | 0.334 |
| EN06 | 0.18 | ± 0.49 |  |  |  |  |
| EN13 | 0.26 | ± 0.34 |  |  |  |  |

Fast Spiking Neurons Firing Rates
| Means and Standard Deviation |  |  |  |  | ANOVA |  |  | Post-hoc test |  |  |  |  |  |  |
| --- | --- | --- | --- | --- | --- | --- | --- | --- | --- | --- | --- | --- | --- | --- |
|  | Stationary |  | Moving |  | Effect | F | p |  | OS | OM | 6S | 6M | 13S | 13M |
| Open | 0.45 ± | 0.47 | 0.75 ± | 0.53 | Group | 0.30 | 0.740 | OS |  |  |  |  |  |  |
| EN06 | 0.54 ± | 0.45 | 0.77 ± | 0.58 | Move | 6.20 | 0.014 | OM |  |  |  |  |  |  |
| EN13 | 0.55 ± | 0.58 | 0.63 ± | 0.77 | G*M | 0.47 | 0.624 | 6S |  |  |  |  |  |  |
|  |  |  |  |  |  |  |  | 6M |  |  |  |  |  |  |
|  |  |  |  |  |  |  |  | 13S |  |  |  |  |  |  |
|  |  |  |  |  |  |  |  | 13M |  |  |  |  |  |  |

**Table 1. Single-Unit firing-rates following enucleation.**
| Means and Standard Deviations |  |  |  | ANOVA |  |  |
| --- | --- | --- | --- | --- | --- | --- |
|  | Move-Stationary |  |  | Effect | F | p |
| Open | 0.30 | ± 0.27 |  | Group | 2.32 | 0.104 |
| EN06 | 0.24 | ± 0.37 |  |  |  |  |
| EN13 | 0.08 | ± 0.39 |  |  |  |  |

To determine if enucleation changed neuronal behavior, we sorted spikes into presumptive single-units. We divided the isolated units into regular-spiking (presumed excitatory) and fast-spiking (presumed inhibitory) clusters and assigned each to superficial (L2-4) or deep layers (L5-6). Firing-rate distributions and means were compared for each group (Fig. 2B & C). An initial 3-factor ANOVA considering Group, Movement, and Layer showed no significant effect for layer, nor any interaction with layer, so layer was not considered further. For regular-spiking neurons, ANOVA revealed significant effects for both Group and Movement but no interaction (Table 1). The group effect results from a decrease in mean firing rates for both EN6 and EN13 during both Stationary and Moving conditions when combined. However, post-hoc tests did not show a significant difference between sham and experimental conditions for the separate Stationary or Moving conditions. Post-hoc analysis confirmed that movement was associated with an increase in firing-rates for all experimental conditions. The amplitude and distribution of movement modulation was not significantly different for regular spiking neurons in either EN6 or EN13. Fast-spiking neurons showed a significant effect of movement by 2-factor ANOVA, but no effect of group or interaction between group and movement. Similarly, the degree of movement modulation was not significantly different for each of the groups. These results show that enucleation (either at P6 or P13) causes a moderate reduction in fast-spiking neuron firing rates across behavioral states, but does not affect the regulation of firing by movement. They confirm results in animals in which pupil dilation can be directly monitored to measure arousal separately from movement (Vinck et al., 2015) that the modulation of the dEEG and the increased firing rates caused by arousal are dissociable because enucleation only strongly affected the dEEG without large changes in firing rates dynamics.

### Role of pattern visual experience in cortical dynamics

Enucleation eliminates the retina, depriving the visual system of both visual experience and spontaneous activity critical for circuit formation and maintenance (Hooks and Chen, 2020). To examine the effects of visual, rather than retinal, deprivation on the establishment of cortical dynamics and to more closely model reversible pattern vision loss by cataracts, we bilaterally sutured the eyelids before natural eye-opening (Sutured group, n=10), and compared them to littermates undergoing a Sham surgery (n=7) on the same day. Eye closure was maintained through the recording period (i.e. the animals were recorded with the eyes sutured) to directly compare with the blind and with the enucleation experiments. For the eye-suturing experiments the animals faced a monitor placed before the contralateral eye (Fig. 3A). Visual stimulation consisted of 1 min blocks of a black screen to test spontaneous activity in the dark, a gray screen to examine low-contrast luminance responses (a visual stimulus likely perceived similarly, though with reduced luminance, by the two groups), and an equiluminant ‘noise’ stimulus consisting of randomly shifting changes in luminance of large squares which should have reduced visibility to the Sutured animals.

**Figure 3.**
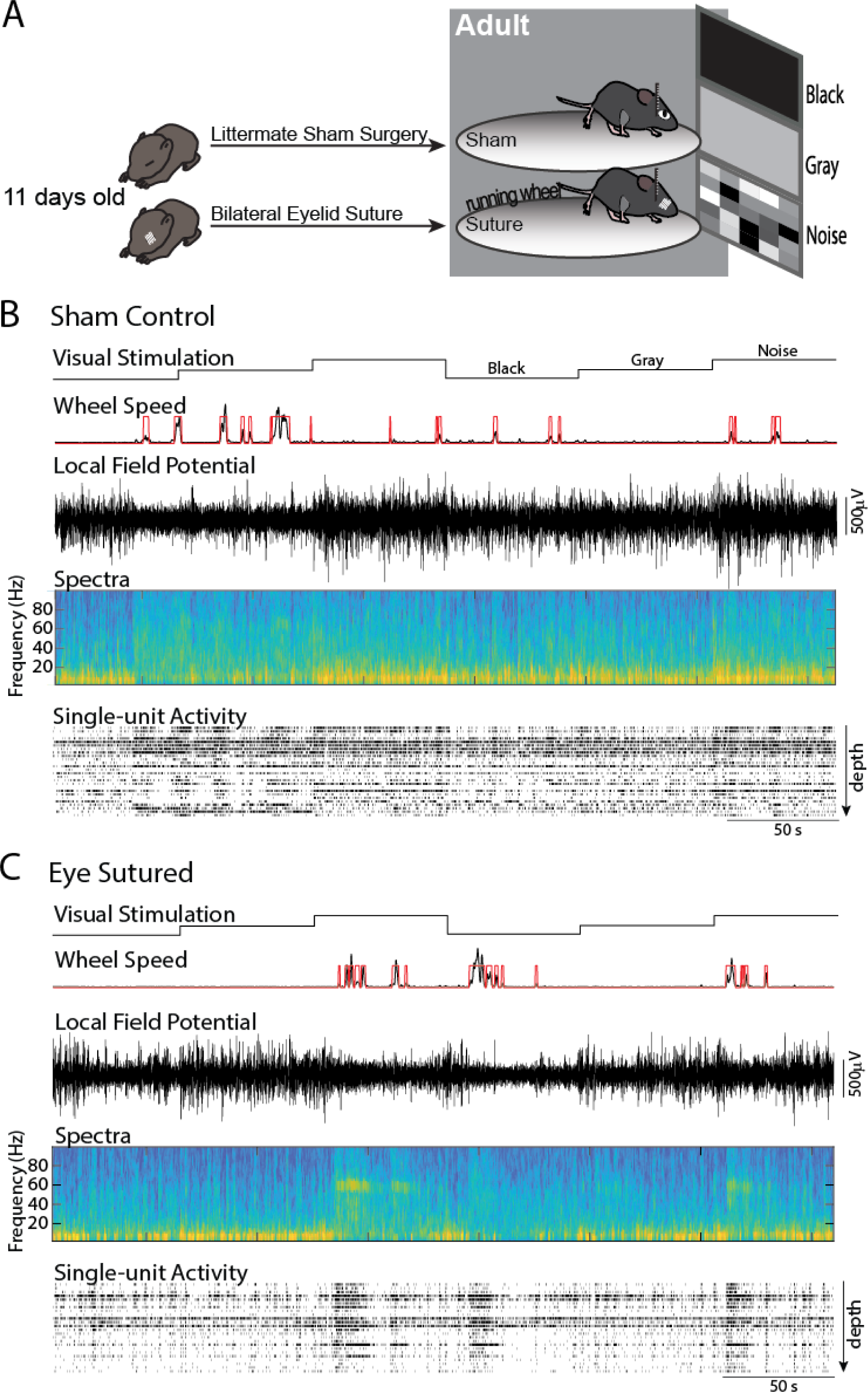
Effect of bilateral eyelid suture on cortical dynamics. **A.** Experimental design. Mouse pups underwent bilateral eyelid suture before natural eye-opening. Control littermates underwent sham surgery and were recorded on alternating days. Activity in the monocular visual cortex was recorded in adult head-fixed, awake animals allowed free movement on a circular treadmill while alternating blocks of visual stimuli were presented to the contralateral eye. Visual stimuli consisted of 1 min blocks of a ‘Black’ screen (full darkness), ‘Gray’ screen’ or isoluminant ‘Noise’ stimuli of changing luminance blocks. **B.** Representative recording from a sham sutured animal across two rounds of stimulation. **C.** Representative recording from eye sutured littermate. Note qualitative similarity in state regulation between sham and sutured animals, including desynchronization of dEEG and increased firing during movement. Also note the increased narrow band gamma generated by this eye-sutured animal during noise stimulation that was amplified by movement.

Sutured and control animals exhibited similar movement dynamics. We observed no significant differences between groups for the total time spent moving (Sham 11.6+/− 2.5%(SEM), Sutured 9.3+/−1.8%, p=0.53 t-test), mean duration of movement bouts (Sham 1419+/−268 ms, Sutured 1489+/−176 ms, p=0.83 t-test), or speed (Sham 345+/−39 arbitrary units, Sutured 320+/−32, p = 0.63 t-test). Neither the distributions of movement, bout length, or speed were different between groups. Like the enucleated animals, the gross neural behavior in the visual cortex was qualitatively similar in Sutured and Sham control animals (Fig. 3B-C). Visual inspection suggested an increase in firing during Noise blocks in Sham animals that was not present in Sutured littermates, while both groups showed strong increases in firing during moving periods in all stimuli conditions. Sutured animals were very likely to display prominent narrow-band gamma power increases during both Gray and Noise blocks. This narrow-band oscillation was present in fewer than half of Sham animals and only during the Gray screen blocks.

We first quantified the behavior of the superficial dEEG. In the dark, Sutured animals displayed a broad-band increase in frequency power during both Stationary and Moving periods (Fig 4A). This increase was significant for the frequencies 2-28Hz during Stationary and 2-56Hz & 70-78Hz for Moving periods. The modulation of the dEEG power by movement was largely similar between groups, but there was a significant decrease in Sutured animals 14-24Hz.

**Figure 4.**
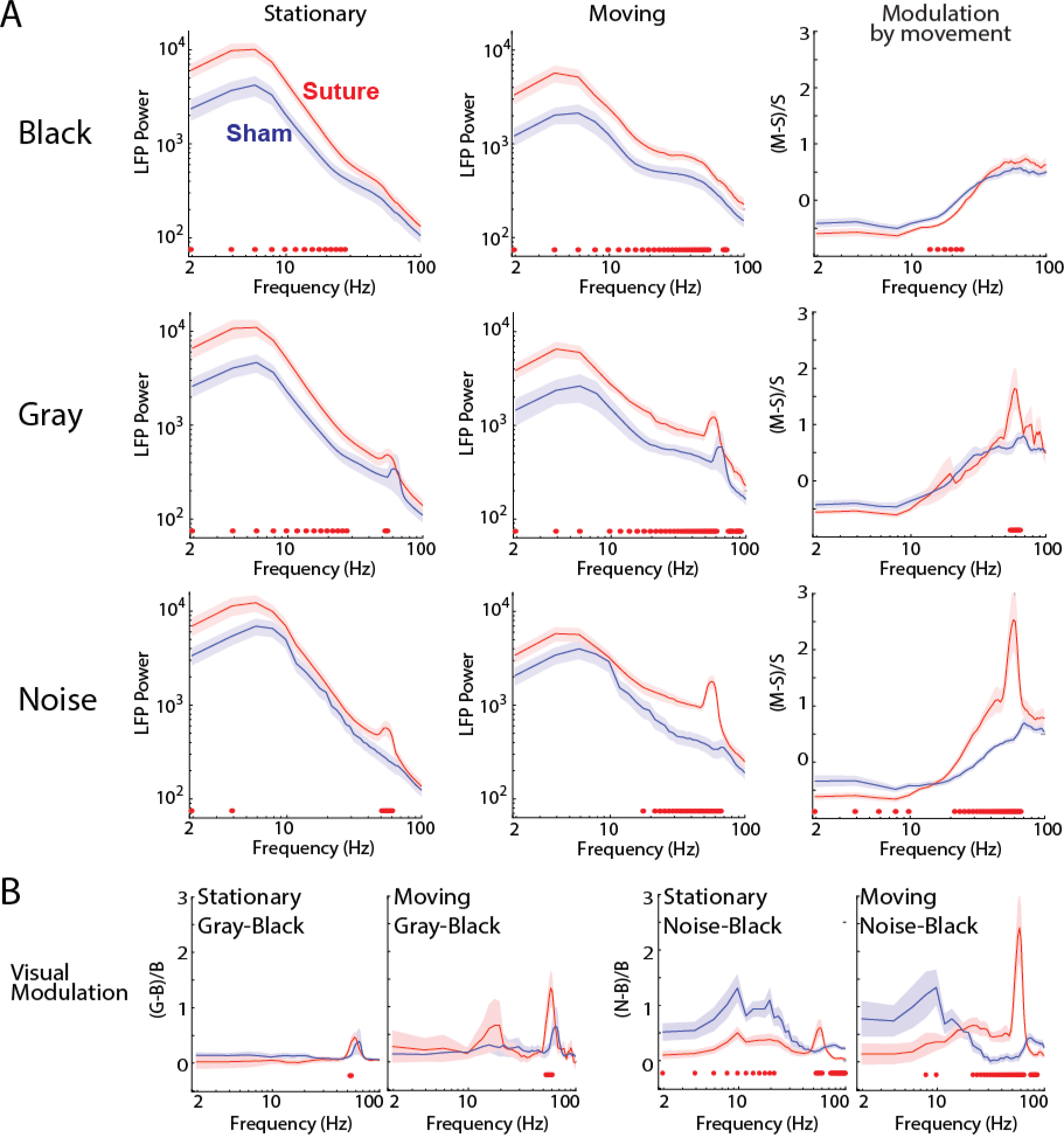
Effect of bilateral eyelid suture on EEG dynamics. **A.** Population mean and standard deviation of dEEG frequency power by visual stimulation (row) and movement state (column). The far right column shows proportional change in power during movement. Frequencies statistically different between groups are marked by purple dots near the x-axis. **B.** Frequency-power change by visual stimulation for each movement condition. Power change for Gray screen(left two) and Noise stimulation(right two) relative to Black screen for Stationary and Moving conditions.

During Gray screen blocks both groups displayed a prominent peak in the spectra resulting from narrowband gamma activity (Storchi et al., 2017; Saleem et al., 2017). The frequency of the peak was shifted to lower frequencies in Sutured animals (peak 64-66Hz in Sham and 56-58Hz in Sutured). As in the Black-screen blocks, overall power was higher for sutured animals 2-28Hz & 52-54Hz during Stationary and 2-60Hz, 74-80Hz & 84-92Hz during Moving. Both groups had a similar decrease in low-frequency and increase in high-frequency dEEG power during movement, but Sutured animals showed a significantly greater increase in the narrowband gamma power(54-64Hz). We quantified the effect of visual stimulation by calculating the proportional change between Black blocks and Gray blocks during Stationary and Moving periods(Fig. 4B). The only significant modulation of dEEG power by Gray-screen was an increase in narrowband gamma power, an increase that was greater during movement. The frequency of narrowband gamma was lower for Sutured animals during both Stationary and Movement periods, while the degree of its modulation was increased specifically during movement.

During Noise blocks the wideband power differences between Sham and Sutured animals were reduced, largely because Sham animals had increased power <20Hz. During Stationary periods the difference between the two groups was limited to the 2-4 & 50-60Hz bands. The later gamma frequency differences reflect the suppression of narrowband gamma in Sham but not Sutured animals, consistent with its origins as a contrast-suppressed luminance activity generated in the retina (Saleem et al., 2017; Storchi et al., 2017). During Movement, Sutured animals had significantly higher power across the broad range of beta-gamma frequencies (18-60Hz). This shift was also reflected in an increased modulation by movement during noise: Sutured animals showed increased suppression of low frequencies (2-10Hz) and a greater increase of high-frequencies (22-66 Hz) during Movement. As suggested by the raw spectra, the effect of patterned visual stimulation was very different between groups. Compared to the frequency powers present in the dark (black screen), Sham animals showed large power increases below 20Hz and above 60Hz during noise stimuli (Fig. 4B right) during both Stationary and Moving periods (significance points not shown). In contrast, these stimulus-driven increases in low and very high frequencies were moderate or absent in the Sutured animals (reduced modulation 2-20 & 72-100Hz Stationary and 8-10 & 74-86Hz Moving). Instead, Sutured animals showed a greater increase in stimulus-driven narrow-band gamma power (52-60Hz) while Stationary, and narrow-band gamma plus lower beta-gamma powers (22-64Hz) while Moving. Overall, the dEEG data show a broadband increase in dEEG power, and normal regulation of the oscillatory regime by state in sutured animals. In addition, sutured animals appear to have increased sensitivity or capacity to generate narrowband gamma oscillations in response to low-contrast luminance. Not surprisingly, Sutured animals generate narrowband gamma activity in response to high-contrast noise stimuli, as this stimulus is filtered by the sutured eyelids, while Sham animals show suppression of narrowband gamma. The origin of the increase in low-frequency activity by Noise stimuli in sighted animals is harder to understand but is likely a combination of the update rate (5 Hz) and intrinsic dynamics. This is lower in the Sutured animals as they have reduced luminance and contrast on the retina.

Neuronal firing rates were also altered by chronic eye closure, with stronger effects observed for regular spiking neurons(Fig. 5 & 6; Table 2 & 3). Population means for firing rates were initially analyzed by 4-factor ANOVA (experimental group vs. move condition vs. stimulation block vs. neuron layer) for RS and FS neurons separately. This analysis showed no significant effect of layer for RS and a moderate effect for FS but no interaction with Group. Layer was therefore not considered further. RS neurons showed significant effects of Group, Movement, and Stimulation, and an interaction between Group and Movement. Examination of the post-hoc tests and firing-rate distributions suggest that these differences result from a decrease in RS firing-rates in Sutured animals that was specific to Stationary periods in all stimulus conditions (Fig. 5A left). Interestingly, firing rates became similar during Moving periods, again for all stimulus conditions (Fig. 5A center column). This normalization of firing rates during movement implies that firing rate increases occurring during movement should be greater in Sutured animals(Fig. 5A right column). 2-factor ANOVA for group and stimulation block on the effect of movement-modulation revealed an effect of group, but not an effect of stimulus block or interaction of the two. In all stimulus conditions sutured animals had larger mean firing rate increases during movement, but this was significant only for the dark blocks.

**Figure 5.**
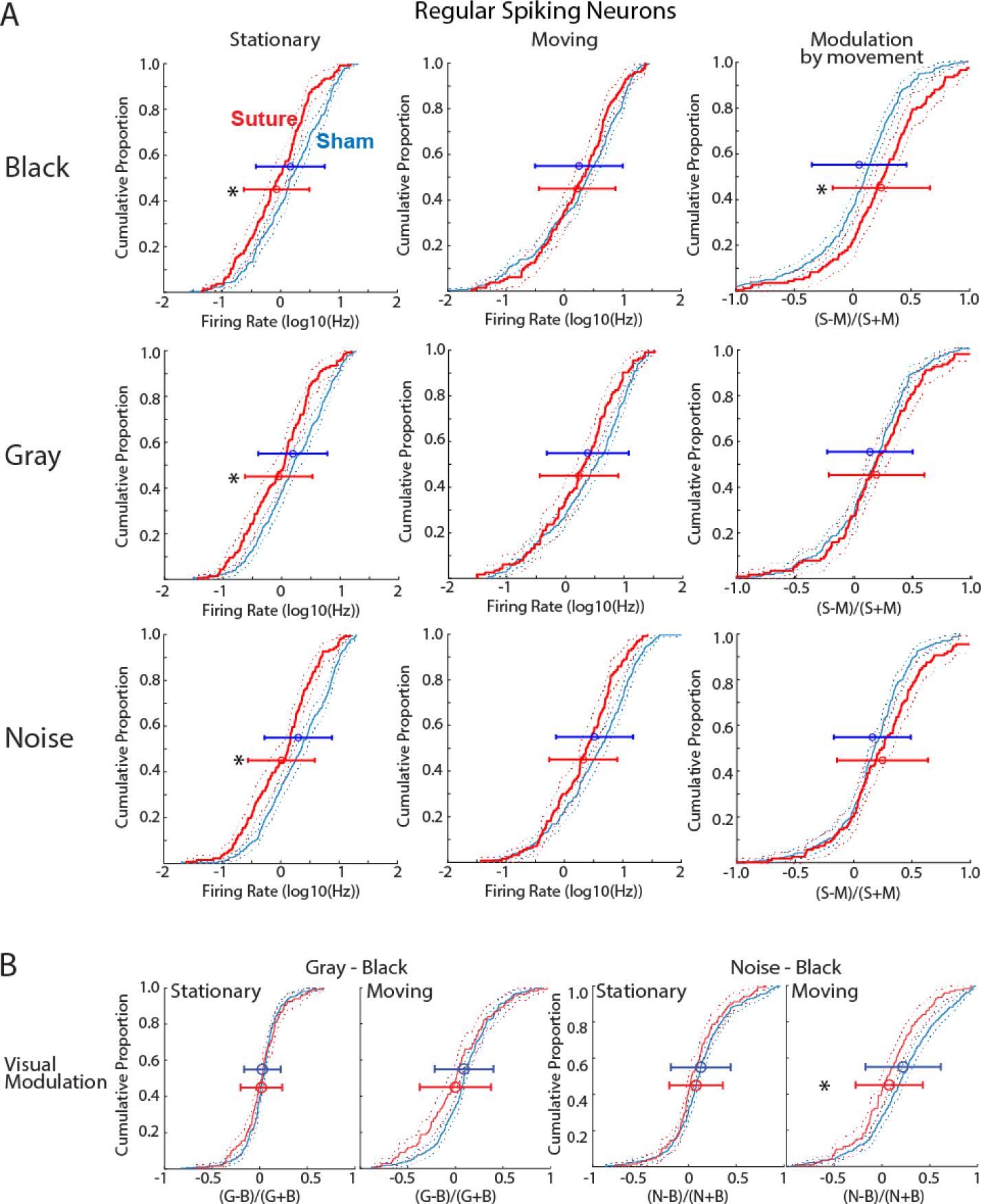
Effect of bilateral eyelid suture on regular spiking neuron firing-rates. **A.** Cumulative distribution histogram of regular spiking neuron firing rates by visual stimulus(row) and movement state(column). Change in firing rate by movement is shown in the right column. Solid lines show distribution and dotted lines the 95% confidence interval. Mean and standard deviation of each distribution is shown by the error bars. Significant difference (ANOVA post-hoc) between groups is shown by asterisk. Firing rates are lower for all stimulus conditions when the animals are stationary, but similar during movement. A significant difference for modulation by movement was observed only in the dark condition. **B.** Cumulative distribution of firing-rate modulation by visual stimulus, divided by behavioral state. Firing rate change relative to black screen is shown for gray screen (left two) and noise(right two) for each Stationary and Moving conditions. See Table 2 for associated data.

**Figure 6.**
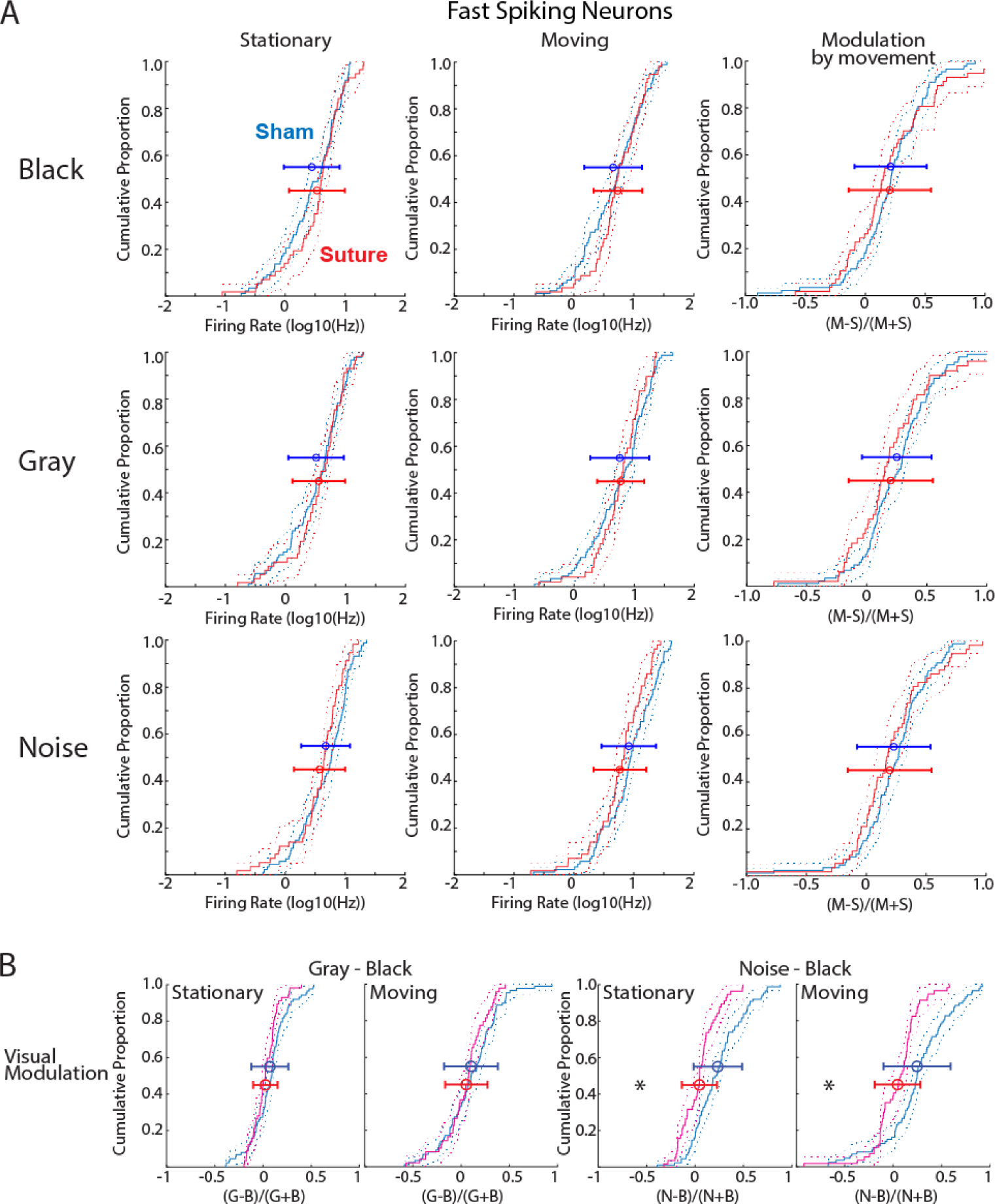
Effect of bilateral eyelid suture on fast spiking neuron firing-rates. **A.** Cumulative distribution histogram of fast-spiking neuron firing rates by visual stimulus(row) and movement state (column). Change in firing rate by movement is shown in the right column. Solid lines show distribution and dotted lines the 95% confidence interval. Mean and standard deviation of each distribution is shown by the error bars. No significant differences were found between groups for any condition. **B.** Cumulative distribution of firing-rate modulation by visual stimulus, divided by behavioral state. Firing rate change relative to black screen is shown for gray screen (left two) and noise(right two) for each Stationary and Moving conditions. Significant difference (ANOVA post-hoc) between groups is shown by asterisk. See Table 3 for associated data.

**Table 2.**
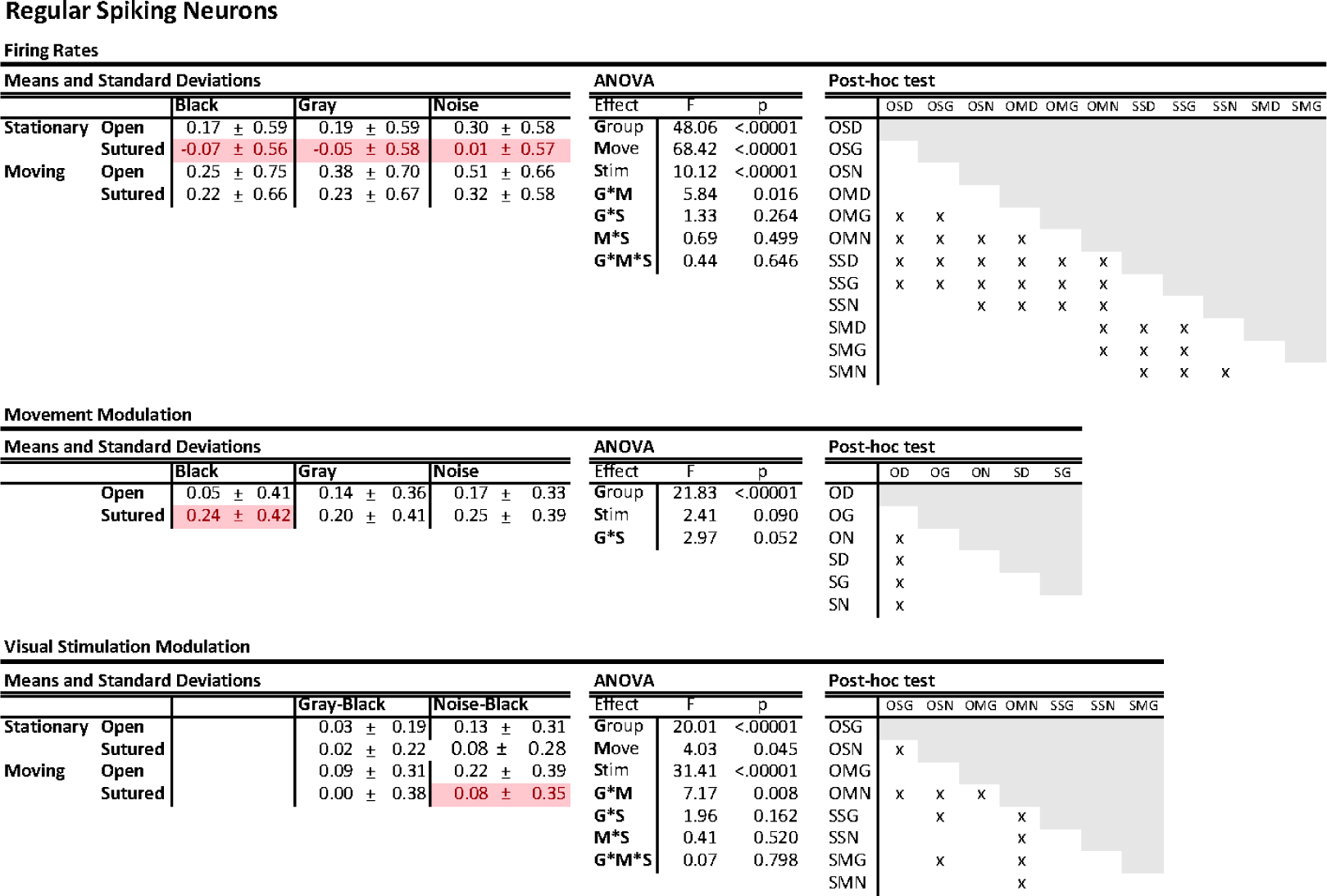
Single-Unit regular-spiking neuron firing-rates following eyelid suture. Means and standard deviations, two-factor ANOVA and post-hoc test results for data displayed in **Fig. 5**. “Open” designates the Sham sutured animals. For means, red indicates significant difference from sham group (by Post-hoc test). For the ANOVA **\*** shows the interaction between factors (eg. **G*M** is the Group **\*** Movement interaction). The post-hoc test results show p<0.05 by Tukey HSD. For the top and bottom charts the first letter (O/S) denotes treatment group; the second letter S/M denotes movement state (**S**tationary/**M**oving), the third letter D/G/N denotes stimulus block (Dark/Gray/Noise).

**Table 3.**
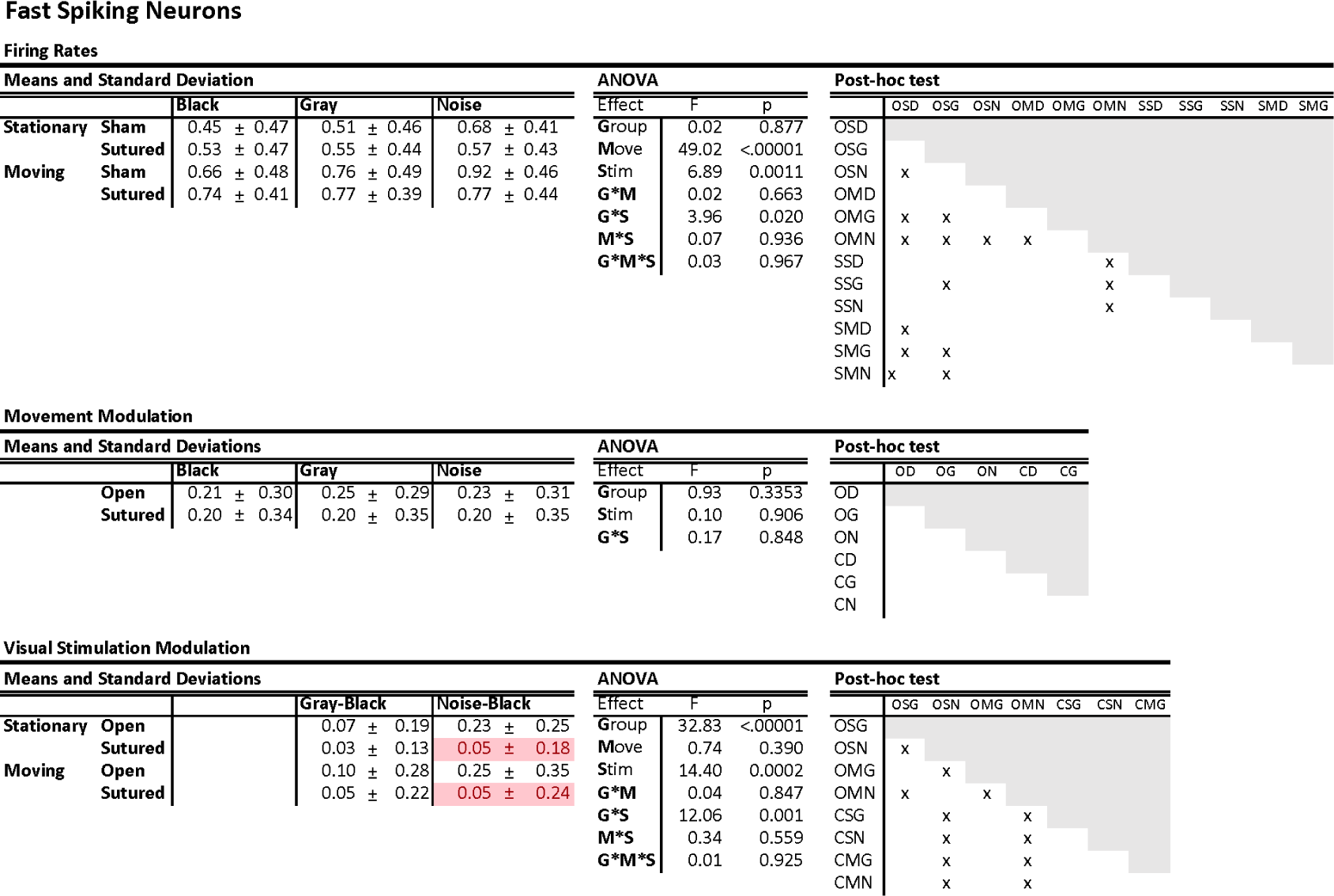
Single-Unit fast-spiking neuron firing-rates following eyelid suture. Means and standard deviations, two-factor ANOVA and post-hoc test results for data displayed in **Fig. 6**. Abbreviations are the same as Table 2.

Examination of firing rate as a function of stimulus condition showed strong increases in firing rates driven by noise, but not gray screen in the sham-treated animals(Fig. 5B). This increased firing by noise stimulus was not present in the Sutured animals. As a result, the firing-rate modulation index for noise stimulation was significantly lower in Sutured animals during both Stationary and moving periods, likely reflecting the lack of patterned vision through their closed eyelids.

The behavior of fast-spiking, presumptive interneurons, was not strongly affected by eye closure. Mean firing rates and distributions were similar between experimental groups for all behavioral and stimulation conditions(Fig. 6A; Table 2). Likewise, movement modulation was similar in both groups. As was observed for regular spiking neurons, neuron firing rates increased during noise stimuli only for Sham animals, and therefore the modulation by noise stimulus was significantly lower for the Sutured animals, likely reflecting their lack of patterned vision (Fig. 6B).

Overall our results confirm that eyelid suture obscures pattern vision but does not impede luminance detection, as reflected in a loss of dEEG and spiking responses to the noise stimulus condition, but increased dEEG narrowband gamma in Sutured animals presented with any visual stimulus despite the attenuation of the closed eyelid. Interestingly the effects of this pattern vision deprivation do not match that of enucleation. Thus chronic eyelid suturing resulted in a reduction in excitatory neuron firing rates during periods of quiet wakefulness and an increase in broad-band dEGG power that was not due to a simple reduction of retinal drive.

### Role of visual and retinal experience in establishing higher-order neuron behavior

Beyond firing rates, we quantified the regularity of neuron firing and the pairwise neural firing rate correlations, parameters that have been shown to differentiate cortical regions and be homeostatically regulated (Hengen et al., 2013; Mochizuki et al., 2016; Wu et al., 2020). To quantify firing patterns we adopted a metric of variance for interspike-interval (LvR) that is independent of firing rate and reveals differences in firing patterns between primate cortical regions (Shinomoto et al., 2009). Regular firing generates a low (<1) LvR, and bursty neurons a high(2+) LvR. Enucleation at P6 and P13 altered the regularity of neuron firing; neurons became more regular than Sham controls (Sham 1.31± 0.30(StDev), EN6 1.25±0.32, EN13 1.20±0.31; ANOVA F=5.46 p=0.004) (Fig. 7A). Eye-suturing, however, did not significantly change the distribution or mean regularity of neurons(1.35±0.31 vs 1.32±0.36, t-test p = 0.32) (Fig. 7B).

**Figure 7.**
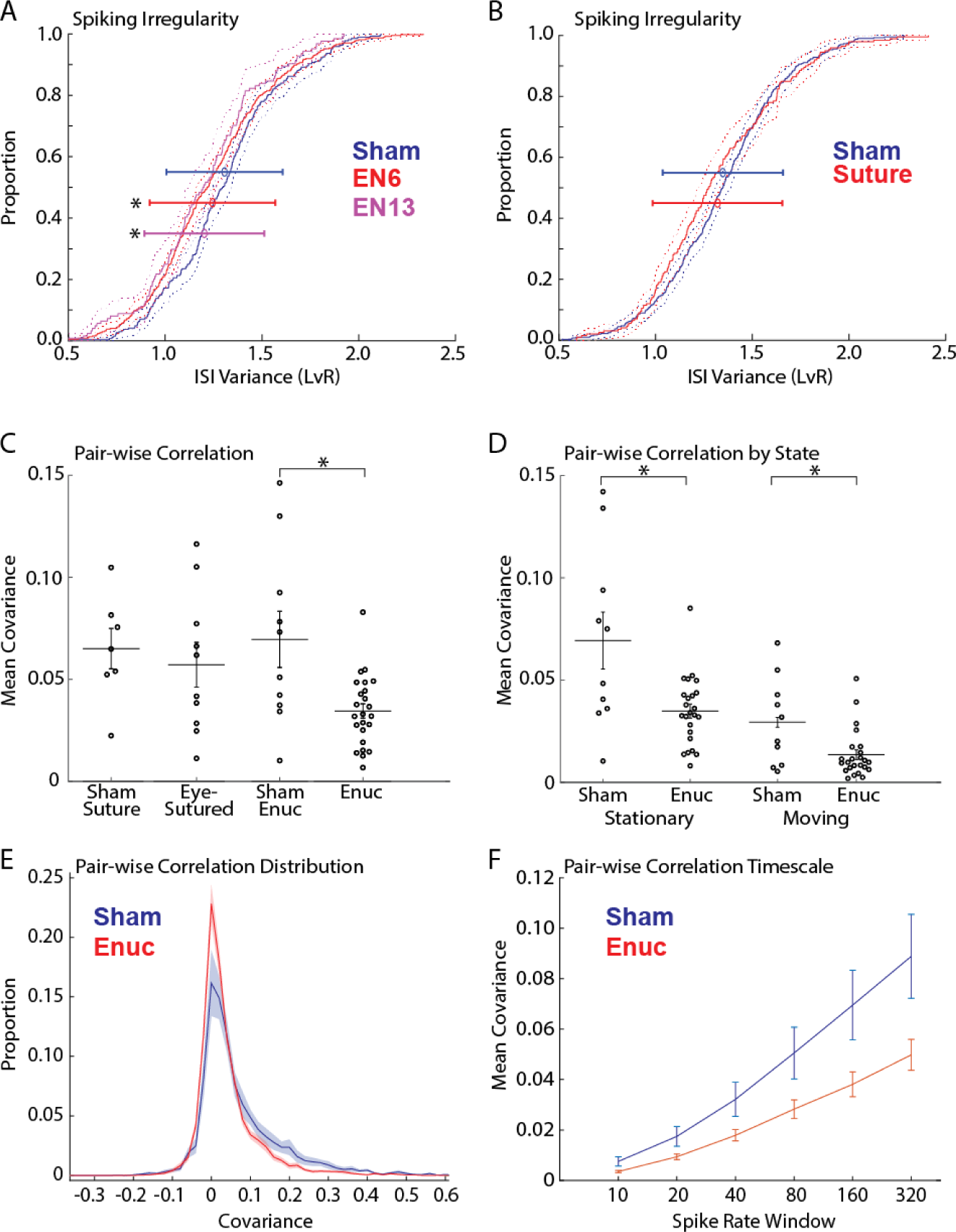
Effect of enucleation and eyelid suture on higher-order firing behavior. **A.** Cumulative distributions of spiking irregularity for the enucleation experiment. LvR is a normalized measure of interspike interval(ISI) that is not affected by firing-rate(Shinomoto et al., 2009); higher LvR indicates less regular firing. Solid lines show distribution, dotted lines 95% confidence interval. Error bars show mean and standard deviation for each group. Significant difference from the Sham group(ANOVA post-hoc) is shown by asterisk. **B.** Cumulative distributions of spiking irregularity for the suturing experiment. **C.** Animal mean pairwise spike rate covariance for both experiments. Mean and standard deviation are shown along with each animal mean covariance measured for a 160 ms window. Asterisk shows significant difference from group control by ANOVA post-hoc. Both P6 and P13 enucleation groups were pooled. **D.** Population mean pairwise spike rate for enucleated groups for Stationary and Moving periods separately(160ms window). Asterisk shows significant difference from group control by ANOVA post-hoc. Reduced correlations caused by enucleation are present for both movement and Stationary periods. **E.** Distribution of pairwise correlations for all neurons(160ms window). Mean and standard deviation(shaded region) shown. **F.** Mean pairwise correlation by window size.

Visual deprivation disrupts the refinement of horizontal connections that link cortical neurons with similar receptive fields (Huberman et al., 2008). However, this shift occurs as a rearrangement of neuronal connections, largely keeping the connection probabilities constant (Ko et al., 2013). To determine if local connections and the resultant correlation between neuron pairs are changed following deprivation, or if it is under long-term homeostatic control as observed for short-term correlation during juvenile development (Wu et al., 2020), we calculated pairwise firing-rate covariance for all the single units isolated in individual animals. The mean pairwise correlation for each animal was then used to compare the experimental groups to their respective controls. As we did not observe any differences between EN6 and EN13 we combined these groups for the analysis. Mean pairwise covariance was similar between the Sutured group and their Sham controls, but was significantly reduced in Enucleated animals compared to their Sham controls (Fig. 7A) (Suture Control 0.065±0.001; Sutured 0.057±0.011; Enucleation Control 0.069±0.014; Enucleated 0.035±0.003) (ANOVA F=4.71; p = 0.0059, with significant differences between the Enucleated and Sham-Enucleated groups only). We asked whether the correlation difference in the enucleated animals was specific to the Stationary or Moving period (Fig 7B). Mean pairwise correlations showed a significant effect for both experimental group and state but no interaction between the two (Sham Stationary 0.069±0.014; En Stationary 0.035±0.003; Sham Moving 0.029±0.007; En Moving 0.013±0.002; 2-factor ANOVA F=18.4, p = 0.0001 for group; F=27.14 p = 0.000002 for state; F= 2.5 p = 0.1185 for interaction). Post-hoc tests show significant mean differences between control and experimental groups for both Stationary and Moving conditions. Thus lower correlations may result from underlying loss of connections rather than changes to the dynamics of either state. Finally, to determine if the lower correlation in enucleated animals is specific to the time window used for correlation calculation, we calculated correlations for multiple windows from 20 to 320 ms (Fig. 7C). We observed lower correlations for all window sizes and these increased with the size of the window (ANOVA for effect of group: F=31.46 p = 6.79*10-8; effect of window size F=31.76 p = 1.09*10-23; interaction F=2.47 p=0.033), indicating no specificity for time-scale in the correlation drop.

Our results show a more profound effect of enucleation on higher-order behavior of neurons, increasing the regularity and decreasing pairwise correlations. In contrast, visual deprivation did not change these behaviors even though it caused a reduction in neuron firing rates that was not observed following Enucleation.

### Role of retinal experience in the development of 3-6 Hz rhythms

Mice frequently display a prominent slow oscillation immediately upon cessation of movement (Einstein et al., 2017). This so-called ‘3-6Hz oscillation’ shares similarities in the pattern of thalamic and cortical engagement with the primate alpha rhythm (Senzai et al., 2019; Nestvogel and McCormick, 2022). Our initial visual inspection of enucleated and sutured animals indicated no loss of the 3-6 Hz oscillation; rather the rhythm appeared more prominent particularly in enucleated animals. Such an increase might be expected if corticothalamic connections are strengthened by enucleation as they are in mice with disrupted retinal development (Seabrook et al., 2013). In our set-up, the peak in dEEG frequency power for this rhythm was closer to 5-8 Hz which largely reflected the frequencies in the individual dEEG spikes, rather than the inter-trough intervals, which were closer to the 3-6Hz described by others. We did not observe a significant change in these frequencies for the non-moving conditions in enucleated animals, suggesting no major regulation of the 3-6Hz pattern. However, any changes may have been masked by the long Stationary periods that did not show the oscillation. In sutured animals, we did observe an increase in power at the relevant frequencies, but this increase occurred as part of a wide-band increase in frequency power that was also present during movement.

To directly test the effects of enucleation on the 3-6Hz oscillation we isolated periods of this oscillation from the remainder of the Stationary periods. The spectra of neither the EN06 nor EN13 groups showed any significant differences of frequency power from control for either the isolated periods of 3-6Hz oscillation or for the remainder of the Stationary periods (Fig. 8A). However, enucleated animals did have an increase in the occurrence of these oscillations (Fig 8B). Periods of 3-6Hz oscillation occupied 2.14±0.39%(SEM) of the recording for Sham animals, 3.05±0.39% in E6 animals and 4.19±0.47% in E13 animals (one-way ANOVA for group F=4.5 p=0.013, with sham significantly different from E13 by post-hoc test). Sutured animals, in contrast, showed increased frequency power during 3-6Hz episodes (significant bands: 2-10, 14-15, 18-20, 24, 28-29Hz). Interestingly, removing the 3-6Hz periods eliminated the significant differences in the spectra for the remainder of the Stationary period (Fig 8C). There were no differences between groups in the incidence of 3-6Hz oscillations (Sham 2.51±0.40% vs Sutured 2.13±0.40%, t-test F-0.42 p=0.53) (Fig. 8D).

**Figure 8.**
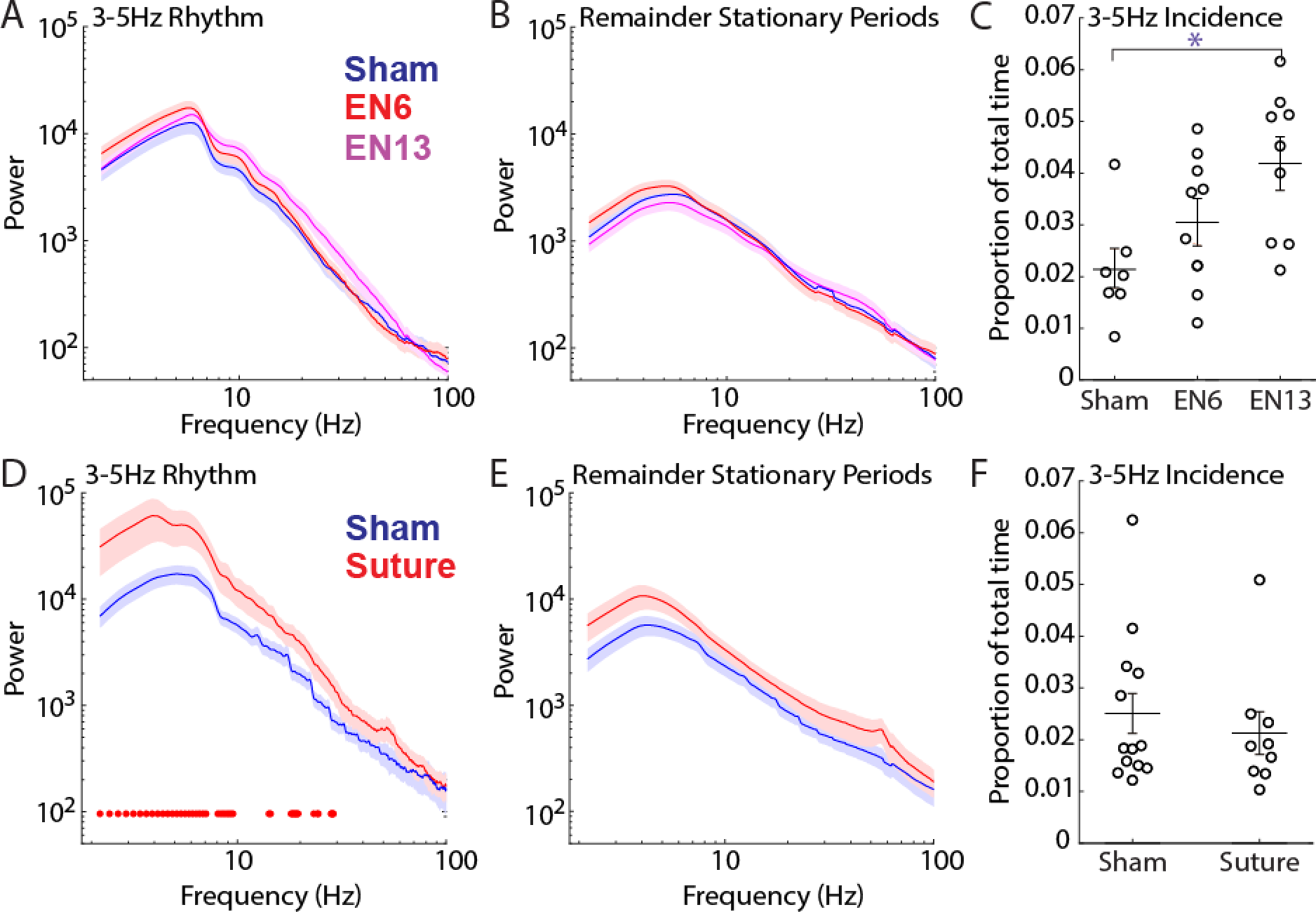
Effect of enucleation and eyelid suture on the 3-6 Hz rhythm. **A.** dEEG frequency power for isolated periods of 3-6 Hz rhythm for the enucleation experiments. Population mean and standard deviation(shaded regions) are shown. No significant differences were observed for any frequency. **B.** dEEG power for the remainder of Stationary periods after removal of 3-6Hz periods. **C.** Proportion of time spent in 3-6Hz rhythm for each animal and the population mean and standard deviation. Significant difference(ANOVA post-hoc) is shown by asterisk. **D-F.** As for A-C with suture experiment. Significant difference between specific frequencies is shown by red dots near x-axis in D.

In sum, our data show that the development of the large amplitude 3-6Hz rhythms depends neither on retinal input nor patterned visual input to the thalamus. In fact, disruption of these inputs increases the incidence or amplitude of 3-6Hz oscillations depending on the form of the deprivation.

### Enucleation identifies differential generative mechanisms of state-dependent low-frequency oscillations

The 3-6 Hz oscillations have been suggested to be a mouse primitive of human alpha oscillations (Senzai et al., 2019). However neither their incidence nor power was reduced by deprivation as is occipital alpha following blindness (Lubinus et al., 2021). The only low-frequency activity reduced by deprivation was the low-frequency activity remaining during movement, which was strongly reduced following enucleation. This suggests that this activity, not the 3-6 Hz oscillations, is a homolog of human alpha. While the spectral profiles of low-frequency activity present during movement and quiescence (both 3-6Hz oscillations and not) are similar, these frequencies could be generated differently, and their generators differentially regulated by visual experience. To test this hypothesis we isolated large amplitude slow-events across the recordings to compare their laminar profiles during the different states. Events were defined as peaks in the band-pass filtered (2-12Hz) LFP. We examined the largest third of events, first assigning them to one of three groups based on their time of occurrence: (1) 3-6Hz periods of high synchronization defined as for the previous section, (2) the remainder of the stationary periods and (3) moving (there were no 3-6 Hz periods during movement). We examined events isolated from L1, L4, and L6 peaks and troughs, but initial observations suggested that the diversity of events could be described from L1 positive peaks and L4 negative peaks, the analysis of which is presented here. We found that in both Sham and Enucleated animals, events isolated during 3-6 Hz periods were similar. They consisted of a prominent L2-5 negativity preceding a large L1 positive deflection and L5-6 negativity (Fig. 9A). Events occurring during movement, however, were different between Sham and Enucleated animals. In Sham animals, only the initial negativity remained and this negativity extended into L1 & L6. Events from Enucleated animals however resembled those isolated during 3-6 Hz periods, though they were smaller. Events from the Sham and Enucleated non-3-6 Hz quiescent period were intermediate between the other states, though closer to those isolated from moving periods.

**Figure 9.**
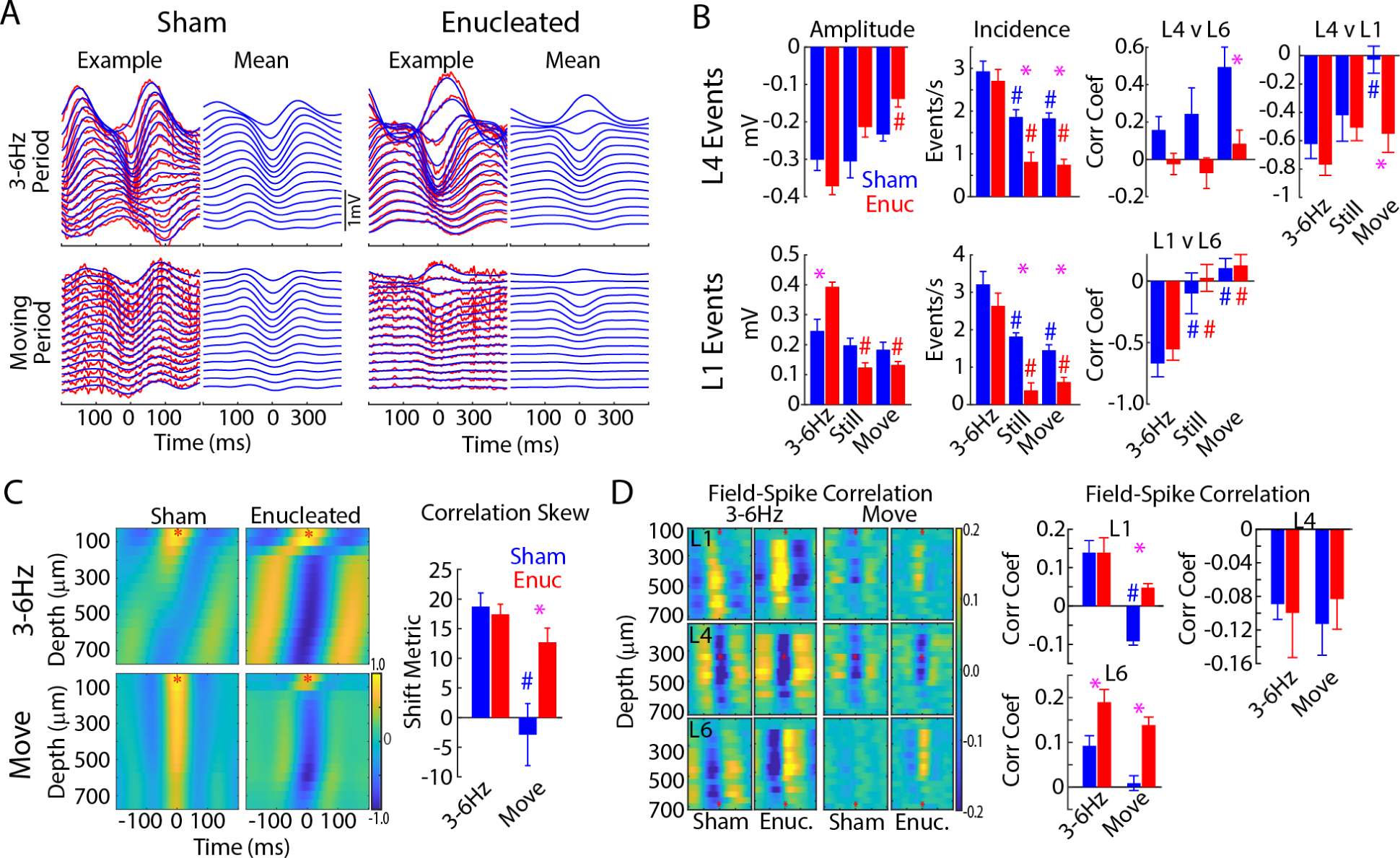
Dissociation of ‘Alpha’ components in Sham and Enucleated animals. **A.** LFP traces through the depth of V1 from representative animals. Left column in each group: A single low-frequency event isolated from the 3-6 Hz Period (top) and Moving Period (bottom). The raw trace is red, and the 2-12 Hz filtered dEEG trace used to isolate and analyze low-frequency events is blue. Right column: the mean traces for events from this animal during each period. Note prominent positive surface peaks and negative deep trough observed during 3-6 Hz but not Moving periods for the Sham animal, while the Enucleated animal has these peaks in both states. **B.** Quantification of low-frequency events identified from negative potentials in L4 (top) or positive potentials in L1 (bottom) during 3-6 Hz periods, quiescent periods without 3-6Hz oscillations (Still), and Moving periods. Population means (n=10 Sham blue, 16 Enuc red) for event Amplitude (left) and Incidence (middle). For this and all graphs, the asterisk denotes the difference between groups for each state, # denotes the difference from the 3-6 Hz period for that group. Incidence but not amplitude of events is significantly lower during Still and Move periods in Enucleated animals compared to Sham, suggesting a reduced number of low-frequency events underlies the reduced frequency power in moving Enucleated animals. The correlation of amplitude in each layer is shown on the right. Correlation analyis shows that the relationship between LFP amplitude in each layer is modulated by state in Sham animals, the correlations between L4 and both L1 and L6 remain constant, suggesting that Enucleated animals lose an event type present during movement in Sham animals. **C.** Representative example of the field-field cross-correlation for each state. Asterisk marks the L1 channel used as the target. (Right) Population means of the difference in the time of peak negativity between adjacent layers on the cross correlogram (‘Correlation Skew’). Sham animals show strong skew for 3-6Hz activity, indicating a spreading wave during these events that is absent during movement. Enucleated animals show similar skew duirng both periods suggesting similar activity patterns. **D.** Analysis of field-spike correlation. (Left) Representative heat-map of the correlation of multiunit firing rate to 2-12 Hz LFP for each period. The LFP layer used for correlation is shown by an asterisk (L1 top, L4 mid, L6 bottom). (Right) Population means of the mean zero-phase field-spike correlation coefficients measured across all layers. Sham animals have different relationships between fields and spikes during 3-6 Hz periods and Moving periods, but while Enucleated animals largely maintain the 3-6 Hz relationship during Moving periods.

To quantify the differences between groups (Sham vs EN) and state (3-6Hz Still, non 3-6Hz Still, Moving), we examined event amplitude and incidence for L1 and L4 derived events (Fig. 9B). Both L1 and L4 events showed significant modulation of amplitudes by state and group with an interaction between state and group caused by larger amplitude reduction in the EN group (L4 two-way ANOVA State F=15.7 p <10-5, Group F=3.08 p=0.083, State*Group F=6.23 p=0.003; L1 F=41.08 p <10-5, Group F=.018 p=0.674, State*Group F=17.54 p <10-5). However, post-hoc analysis revealed no significant difference between Sham and EN for amplitude during movement, but did show an increase in the size of L1 events during 3-6Hz oscillations in the EN group. The incidence of L4 and L1 events was strongly modulated by state, an effect that was greater for Enucleated animals (L4 State F=32.19 p <10-5, Group F=19.6 p <10-5, State*Group F=2.5 p=0.089; L1 State F=39.43 p <10-5, Group F=23.06 p <10-5, State*Group F=1.64 p=0.200). Post-hoc analysis reveled a significantly lower incidence of both L1 and L4 events in EN animals both during movement and during the non-3-6Hz Still periods. Together these data suggest the reduction in low-frequency power during movement comes about through a combination of the decreased incidence and amplitude of slow network events, and with EN animals more strongly reducing the incidence of such events.

To further examine the potential for changes in the structure of these network events, we quantified the correlation between LFP amplitude in different layers. The correlation between the peak LFP negativity in L4 and L6 during L4 events was lowered by Enucleation for all states, though this only reached significance during movement periods, as L4 and L6 became increasingly correlated in Sham but not EN animals (L4vL6 amplitude correlation State F=4.19 p <0.019, Group F=18.86 p <10-5, State*Group F=0.91 p=0.406). The correlation between dEEG amplitude in L1 and L4 for L4 events showed an important interaction between state and correlation. In Sham animals L4 amplitudes are negatively correlated to L1 amplitudes in 3-6Hz periods, but not during movement. However, they remained negatively correlated in Enucleated animals (State F=7.14 p=0.001; Group F=3.75 p=0.058; State*Group F=3.96 p=0.023). The correlation between amplitude in L1 and L6 for L1 events was not changed by Enucleation: both groups showed a significant negative correlation during 3-6Hz periods that was eliminated during Still and Movement periods (L1vL6 State F=25.83 p <10-5, Group F=1.00 p=0.320, State*Group F=0.14 p=0.869). In total our event analysis suggests that in Sham animals low-frequency events consist of two dissociable circuit components: a translaminar negativity originating in L4 and a L6-L1 dipole. Together these components result in L1-L6 and L4-L1 anti-correlation during 3-6Hz events but allow for increased interlaminar correlation when the L6-L1 dipole disappears during movement. Thus the 3-6Hz rhythm is composed of the two components, but during movement (and to a lesser extent during the non-3-6Hz quiescent periods) the L4 event alone is prevalent and the combined events are rare. In the EN animals however the L4 events are not generated during movement and only a few, lower amplitude, combined events remain.

Analysis of low-frequency events requires thresholding of the data in a manner that is necessarily arbitrary and may hide differences between groups and states. To analyze the layer interactions in a more unbiased manner we examined the correlation between local dEEG in different layers and between the dEEG and multi-unit spiking for each state. Field-field cross-correlation revealed a strong difference in the temporal organization of activity between 3-6Hz periods and moving periods for Sham, but not Enucleated animals (Fig 9C). The cross-correlation between layers (here L1 presented) during 3-6Hz activity shows a prominent temporal shift as depolarization travels between layers; this shift occurred both in Sham and Enucleated animals. However, during movement, this shift was not apparent in Sham animals and all layers were positively correlated with zero phase offset. Enucleated animals however maintained the temporal shift, though the amplitude of the correlations was reduced (Correlation Skew: two-way ANOVA State F=22.17 p <10^−5^, Group F=6.48 p=0.014, State*Group F=9.14 p=0.004).

Finally, we examined the cross-correlation between LFP and multi-unit firing rates through the depth of V1. As for the other measures this analysis showed a significant shift in the nature of synchronization between 3-6Hz periods and moving periods that was present in the Sham animals, but not Enucleated (Fig 9D). In particular, the relationship between firing – rates and L1 LFP shifted from positive to negative in Sham animals between states but remained positive for Enucleated (two-way ANOVA State F=29.1 p <10^−5^, Group F=5.47 p=0.025, State*Group F=5.41 p=0.026). For L6 during the 3-6 Hz oscillation, we observed a strong negative correlation immediately preceding the zero point and a positive correlation immediately following. Quantifying the positive correlation showed a higher amplitude for Enucleated animals in both states, but a similar behavior between states (two-way ANOVA State F=7.21 p=0.011, Group F=20.67 p=10^−4^, State*Group F=0.42 p=0.521). For L4 both groups were similar with strong negative correlations between L4 LFP and firing throughout the cortical depth (two-way ANOVA State F=0.01 p=0.938, Group F=0.04 p=0.846 State*Group F=0.2 p=0.658).

In total our detailed analysis suggests that low-frequency activity consists of two dissociable components that occur together during 3-6 Hz periods: an ‘active’ L4 component that drives firing throughout much of the cortex and an ‘inactive’ down-state that suppresses activity and results in a large positive amplitude deflection in L1. During movement the second component is suppressed and Sham animals produce synchronized firing using only the activating component. Enucleated, animals appear to be less able to generate the first, active, component during movement resulting in fewer low-frequency events and lower spectral power in these frequencies. The ‘bare’ active component of these low-frequency rhythms is likely to be closer to a mouse homolog of the alpha rhythm as it was reduced by enucleation.

## Discussion

We used extracellular polytrode recordings of ongoing background activity in awake, head-fixed adult mice allowed to run freely on a treadmill to explore how two different modes of chronic input deprivation, pattern vision deprivation with eye-lid suture and retinal enucleation, modify the spontaneous activity dynamics of the primary visual cortex. We find that spiking and dEEG behavior is largely normal in both cases. However, each deprivation caused specific changes that were non-overlapping and not additive. Thus, we show that the establishment and maintenance of cortical network properties, while experience-dependent, requires neither the primary sense organ nor pattern vision. Enucleation altered multiple cortical network properties including cortical oscillations, correlations, and firing patterns, but exerted only moderate effects on firing rates. In contrast, eyelid suture reduced firing rates of excitatory neurons but left other properties largely intact. Neither deprivation reduced the incidence or size of the prominent 3-6 Hz oscillations that occur sporadically following movement, as would be expected if they are a mouse homolog of human alpha as they are reduced by congenital blindness (Hawellek et al., 2013; Lubinus et al., 2021). However, we did observe that enucleation causes a substantial reduction of low-frequency dEEG power specifically during periods of movement. This activity in intact animals is driven by a significantly different pattern of laminar generators than the 3-6Hz oscillations and is specifically lost following enucleation, suggesting that the low-frequency oscillations remaining during movement (and likely general arousal) are the mouse homolog of human alpha.

### Mouse visual ‘alpha’

Low-frequency activity during movement was reduced by enucleation, but not lid suture. This is consistent with reports of occipital alpha loss in congenitally blind individuals, particularly those with severe congenital vision loss (Bottari et al., 2016; Jan and Wong, 1988). Posterior alpha is present during development even in the blind (Campus et al., 2021), suggesting a regressive process disassembles alpha generators in the absence of retinal input. Interestingly the low-frequency loss we observed was specific to movement periods, when these frequency bands are suppressed as part of the activation resulting from arousal and movement (McCormick et al., 2014; Niell and Scanziani, 2021). Based on similarities of the depth profile (Senzai et al., 2019) and visual modulation (Einstein et al., 2017), large amplitude ‘3-6Hz’ rhythms occurring during quiescence have been suggested to be a mouse primitive of alpha. Rather than losing these oscillations, however, as we would expect if blindness has a similar effect in mice and humans, we observed increased post-movement 3-6Hz bursts in both experimental conditions. This is similar to (Reimer et al., 2014) who observed no loss of these oscillations after retinal degeneration. The homology of 3-6Hz oscillations and human alpha is further argued against by their generation by low-threshold spike bursts in thalamus(Nestvogel and McCormick, 2022), which is not a likely mechanism of alpha(Crunelli et al., 2018). Furthermore, the temporal relationship between pupil dilation and low-frequency/alpha power is different in humans and rodents(Montefusco-Siegmund et al., 2022). We instead suggest the 3-6 Hz oscillations are homologs of human delta which shows increased power in blind adults and infants (Campus et al., 2021).

The alpha-band may intermix multiple activities with different cognitive functions and different generative mechanisms(Alamia et al., 2023; Bollimunta et al., 2008; van Kerkoerle et al., 2014). We identified two circuit contributors to presumptive ‘alpha’ in mice, a rhythmic input likely generated in the thalamocortical loop that is present during movement and quiescence (Molnár et al., 2021), and an inhibitory component present only during the 3-6Hz bursts. The inhibitory component could be mediated by increased inhibition or increased synchronization and bursting of the thalamic relay neurons with reduced arousal, or both(Nestvogel and McCormick, 2022). One interpretation of our results is that synchronization of the thalamus by high-threshold spikes is reduced following enucleation, while the low-threshold induced bursting is not. This is expected to cause strong synchronization during low arousal states but reduced ‘alpha’ during arousal(Crunelli et al., 2018). Overall our results suggest disruption of a thalamic oscillator as the mechanism of alpha loss in the blind(Fiebelkorn et al., 2019). By identifying multiple components of the potential alpha oscillations present in awake mice such as their state-dependence and sensitivity to blindness, the present results provide potential avenues to study alpha circuitry in this tractable model system. The fact that we identified ‘mouse alpha’ as limited to movement periods is likely a side-effect of our use of running as the sole monitor of arousal. Cortical LFP oscillations are more closely correlated with more sensitive measures of arousal like pupil dilation, with movement occurring at the end of the scale(McCormick et al., 2020). Given the changes we observed during still activity outside of the 3-6 Hz bursts, it is likely that mouse alpha will also be present during sedentary arousal and behavioral engagement, and is not linked to movement *per se*. While we did not observe large changes in the behavior of enucleated mice, it remains possible that changes in sensory engagement (e.g. more whisking or attention to audition) contribute to the enucleated phenotype. This is also likely true in human blindness. Furthermore, the large majority of movement-related activity shifts were intact and quantitatively similar between groups, arguing against the existence of significant differences between the moving state in enucleated and sham animals.

### Homeostasis and circuit changes in blindness

Single neuron firing-rates as well as higher-order network behaviors are homeostatically regulated in mouse visual cortex, returning to normal levels within days of manipulations that reduce or increase firing(Wen and Turrigiano, 2024). Our results indicate however that homeostatic set-points can be altered by the profound changes in excitatory and inhibitory circuits resulting from visual deprivation (Ribic, 2020). While enucleation at P6--when spontaneous firing rates are 10% of adults (Riyahi et al., 2021)--did not significantly change adult firing rates, binocular deprivation–a procedure that causes less acute reduction in firing-rates and allows for greater homeostatic plasticity in adults (Hengen et al., 2013; Keck et al., 2013)--did lower firing-rates, but only during Stationary periods. It is unclear whether mean firing-rates were truly lower in sutured animals and an increased movement-related signal normalized them, or whether the firing-rate set points are similar in both animals but set during the movement period. Firing-rate ‘upscaling’ following monocular deprivation occurs during wake (Hengen et al., 2016) suggesting this may also be a critical state in determining when the neuron has returned firing rates to the setpoint.

Higher-order network behaviors such as pairwise correlation, firing regularity, and network criticality are also rapidly altered by visual deprivation but recover on different timescales (Hengen et al., 2013; Ma et al., 2019; Wu et al., 2020), likely because they use different plasticity mechanisms (Keck et al., 2017). These different mechanisms were apparent in our data as pairwise correlations and regularity were altered by enucleation despite normal firing-rates, while eyelid-suture reduced firing without changing correlation or regularity. Horizontal connections are selectively formed between neurons with similar visual response properties while maintaining connection probabilities(Ko et al., 2013). Significant numbers of correlated neurons are first observed around P15, after the times of enucleation here (Colonnese et al., 2017). Our data thus suggest that even though the level of correlation and regularity is largely intrinsically determined, it can be developmentally influenced by changes in activity. An open question is whether changes in these network properties are the result of cross-modal input plasticity or the simple absence of retinal input. Spike intervals are more regular in ‘higher-order’ structures (Shinomoto et al., 2009), which receive a larger portion of their input from within the thalamocortical loop, and bilateral enucleation increases thalamocortical and cortico-cortical connections (Karlen et al., 2006). Thus strengthened corticothalamic input may underlie the increased regularity in enucleated animals, making V1 more like ‘higher’ cortex.

### Cortical gamma oscillations and visual experience

Narrowband gamma oscillations are generated in the retina to low-contrast luminance while broadband oscillations iare generated in cortex in response to high-contrast, structured stimuli (Saleem et al., 2017; Veit et al., 2017; Storchi et al., 2017). Together these activities encode contrast and reflect multiplexed visual channels (Meneghetti et al., 2021). We observed an increase in both types following visual deprivation, but not after enucleation. Narrowband gamma occurred at a lower frequency but higher power in sutured animals. Narrowband gamma emerges during the fourth week with a central frequency around 52Hz, and is delayed by dark-rearing (Chen et al., 2015). Our data indicate extended visual deprivation does not prevent development of narrowband gamma, but does prevent the increase in its central frequency with age. Lid-suture likely increases visual sensitivity, as these animals have larger and more prevalent gamma oscillations despite lowered luminance levels as a result of the closed eyelids. This may be the result of decreased inhibition as both enucleation and suture disrupt the development of cortical inhibitory circuits (Ossandón et al., 2023; Ferrer and De Marco Garcia, 2022).

## Conclusions

Our results show that in mice, like humans, the patterns of ongoing network activity in the visual cortex and their regulation by state are homeostatically regulated and largely independent of input. The primary exceptions are the alpha frequency oscillations which are linked to reduced cortical attention during aroused states in visual cortex (Van Diepen et al., 2019). This alpha suppression may contribute to continued visual impairment following blindness reversal (Maurer et al., 2007; Kalia et al., 2014; Röder and Kekunnaya, 2022). By identifying the likely mouse homolog of alpha and identifying the parameters of its experience-dependence, we provide possible methods to study its generation and plasticity in a tractable, genetic model system.

## Acknowledgements

The authors thank Dr Mary Ann Stepp (GWU) for her guidance on corneal and lens health assays and ocular health assessments.

## REFERENCES

Adrian ED, Matthew BHC (1934) The Berger rhythm: Potential changes from the occipital lobes in man. Brain 57:355–385.

Alamia A, Terral L, D’ambra MR, VanRullen R (2023) Distinct roles of forward and backward alpha-band waves in spatial visual attention. eLife 12:e85035.

Bollimunta A, Chen Y, Schroeder CE, Ding M (2008) Neuronal mechanisms of cortical alpha oscillations in awake-behaving macaques. J Neurosci 28:9976–9988.

Bottari D, Troje NF, Ley P, Hense M, Kekunnaya R, Röder B (2016) Sight restoration after congenital blindness does not reinstate alpha oscillatory activity in humans. Sci Rep 6:24683.

Brainard DH (1997) The psychophysics toolbox. Spat Vis 10:433–436.

Burton H (2003) Visual cortex activity in early and late blind people. J Neurosci 23:4005–4011.

Buzsáki G, Draguhn A (2004) Neuronal oscillations in cortical networks. Science 304:1926–1929.

Campus C, Signorini S, Vitali H, De Giorgis V, Papalia G, Morelli F, Gori M (2021) Sensitive period for the plasticity of alpha activity in humans. Dev Cogn Neurosci 49:100965.

Chen G, Rasch MJ, Wang R, Zhang XH (2015) Experience-dependent emergence of beta and gamma band oscillations in the primary visual cortex during the critical period. Sci Rep (England) 5:17847.

Cohen MX (2014) Analyzing neural time series data: Theory and practice. Cambridge, MA: MIT Press.

Colonnese MT (2014) Rapid developmental emergence of stable depolarization during wakefulness by inhibitory balancing of cortical network excitability. J Neurosci 34:5477–5485.

Colonnese MT, Shen J, Murata Y (2017) Uncorrelated neural firing in mouse visual cortex during spontaneous retinal waves. Frontiers in Cellular Neuroscience 11:289.

Crunelli V, Lorincz ML, Connelly WM, David F, Hughes SW, Lambert RC, Leresche N, Errington AC (2018) Dual function of thalamic low-vigilance state oscillations: Rhythm-regulation and plasticity. Nat Rev Neurosci (England) 19:107–118.

Einstein MC, Polack P, Tran DT, Golshani P (2017) Visually evoked 3-5 hz membrane potential oscillations reduce the responsiveness of visual cortex neurons in awake behaving mice. J Neurosci 37:5084–5098.

Ferrer C, De Marco Garcia NV (2022) The role of inhibitory interneurons in circuit assembly and refinement across sensory cortices. Front Neural Circuits (Switzerland) 16:866999.

Fiebelkorn IC, Pinsk MA, Kastner S (2019) The mediodorsal pulvinar coordinates the macaque fronto-parietal network during rhythmic spatial attention. Nat Commun 10:215.

Golding B, Pouchelon G, Bellone C, Murthy S, Di Nardo AA, Govindan S, Ogawa M, Shimogori T, Luscher C, Dayer A, Jabaudon D (2014) Retinal input directs the recruitment of inhibitory interneurons into thalamic visual circuits. Neuron (United States) 81:1057–1069.

Harris KD, Thiele A (2011) Cortical state and attention. Nat Rev Neurosci (England) 12:509–523.

Hawellek DJ, Schepers IM, Roeder B, Engel AK, Siegel M, Hipp JF (2013) Altered intrinsic neuronal interactions in the visual cortex of the blind. J Neurosci 33:17072–17080.

Hengen KB, Torrado Pacheco A, McGregor JN, Van Hooser SD, Turrigiano GG (2016) Neuronal firing rate homeostasis is inhibited by sleep and promoted by wake. Cell (United States) 165:180–191.

Hengen KB, Lambo ME, Van Hooser SD, Katz DB, Turrigiano GG (2013) Firing rate homeostasis in visual cortex of freely behaving rodents. Neuron (United States) 80:335–342.

Hooks BM, Chen C (2020) Circuitry underlying experience-dependent plasticity in the mouse visual system. Neuron (United States) 107:986–987.

Hoy JL, Niell CM (2015) Layer-specific refinement of visual cortex function after eye opening in the awake mouse. J Neurosci (United States) 35:3370–3383.

Huberman AD, Feller MB, Chapman B (2008) Mechanisms underlying development of visual maps and receptive fields. Annu Rev Neurosci 31:479–509.

Jan JE, Wong PK (1988) Behaviour of the alpha rhythm in electroencephalograms of visually impaired children. Dev Med Child Neurol 30:444–450.

Jeavons PM (1964) The electro-encephalogram in blind children. Br J Ophthalmol (England) 48:83–101.

Kalia A, Lesmes LA, Dorr M, Gandhi T, Chatterjee G, Ganesh S, Bex PJ, Sinha P (2014) Development of pattern vision following early and extended blindness. Proc Natl Acad Sci U S A 111:2035–2039.

Kampf-Lassin A, Wei J, Galang J, Prendergast BJ (2011) Experience-independent development of the hamster circadian visual system. PLoS One 6:e16048.

Karlen SJ, Kahn DM, Krubitzer L (2006) Early blindness results in abnormal corticocortical and thalamocortical connections. Neuroscience 142:843–858.

Keck T, Hubener M, Bonhoeffer T (2017) Interactions between synaptic homeostatic mechanisms: An attempt to reconcile BCM theory, synaptic scaling, and changing excitation/inhibition balance. Curr Opin Neurobiol (England) 43:87–93.

Keck T, Keller GB, Jacobsen RI, Eysel UT, Bonhoeffer T, Hubener M (2013) Synaptic scaling and homeostatic plasticity in the mouse visual cortex in vivo. Neuron (United States) 80:327–334.

Kiorpes L (2016) The puzzle of visual development: Behavior and neural limits. J Neurosci (United States) 36:11384–11393.

Ko H, Cossell L, Baragli C, Antolik J, Clopath C, Hofer SB, Mrsic-Flogel TD (2013) The emergence of functional microcircuits in visual cortex. Nature (England) 496:96–100.

Lubinus C, Orpella J, Keitel A, Gudi-Mindermann H, Engel AK, Roeder B, Rimmele JM (2021) Data-driven classification of spectral profiles reveals brain region-specific plasticity in blindness. Cerebral Cortex 31:2505–2522.

Ma Z, Turrigiano GG, Wessel R, Hengen KB (2019) Cortical circuit dynamics are homeostatically tuned to criticality in vivo. Neuron (United States) 104:655–664.e4.

Maurer D, Mondloch CJ, Lewis TL (2007) Effects of early visual deprivation on perceptual and cognitive development. Prog Brain Res (Netherlands) 164:87–104.

McCormick DA, Nestvogel DB, He BJ (2020) Neuromodulation of brain state and behavior. Annu Rev Neurosci (United States) 43:391–415.

McCormick DA, McGinley MJ, Salkoff DB (2014) Brain state dependent activity in the cortex and thalamus. Curr Opin Neurobiol 31C:133–140.

McGinley MJ, Vinck M, Reimer J, Batista-Brito R, Zagha E, Cadwell CR, Tolias AS, Cardin JA, McCormick DA (2015) Waking state: Rapid variations modulate neural and behavioral responses. Neuron (United States) 87:1143–1161.

Meneghetti N, Cerri C, Tantillo E, Vannini E, Caleo M, Mazzoni A (2021) Narrow and broad γ bands process complementary visual information in mouse primary visual cortex. eNeuro 8:.

Merabet LB, Pascual-Leone A (2010) Neural reorganization following sensory loss: The opportunity of change. Nat Rev Neurosci 11:44–52.

Mishina M, Senda M, Kiyosawa M, Ishiwata K, De Volder AG, Nakano H, Toyama H, Oda K, Kimura Y, Ishii K, Sasaki T, Ohyama M, Komaba Y, Kobayashi S, Kitamura S, Katayama Y (2003) Increased regional cerebral blood flow but normal distribution of GABAA receptor in the visual cortex of subjects with early-onset blindness. Neuroimage 19:125–131.

Mitchell DE, Maurer D (2022) Critical periods in vision revisited. Annu Rev Vis Sci 8:291–321.

Mitra P, Bokil H (2007) Observed brain dynamics. New York: Oxford University Press, USA.

Miyamoto H, Katagiri H, Hensch T (2003) Experience-dependent slow-wave sleep development. Nat Neurosci (United States) 6:553–554.

Mochizuki Y et al (2016) Similarity in neuronal firing regimes across mammalian species. J Neurosci (United States) 36:5736–5747.

Molnár B, Sere P, Bordé S, Koós K, Zsigri N, Horváth P, Lőrincz ML (2021) Cell type-specific arousal-dependent modulation of thalamic activity in the lateral geniculate nucleus. Cereb Cortex Commun 2:tgab020.

Montefusco-Siegmund R, Schwalm M, Rosales Jubal E, Devia C, Egaña JI, Maldonado PE (2022) Alpha EEG activity and pupil diameter coupling during inactive wakefulness in humans. eNeuro 9:.

Murata Y, Colonnese MT (2018) Thalamus controls development and expression of arousal states in visual cortex. J Neurosci (United States) 38:8772–8786.

Nestvogel DB, McCormick DA (2022) Visual thalamocortical mechanisms of waking state-dependent activity and alpha oscillations. Neuron 110:120–138.e4.

Niell CM, Scanziani M (2021) How cortical circuits implement cortical computations: Mouse visual cortex as a model. Annu Rev Neurosci 44:517–546.

Niell CM, Stryker MP (2010) Modulation of visual responses by behavioral state in mouse visual cortex. Neuron (United States) 65:472–479.

Noebels JL, Roth WT, Kopell BS (1978) Cortical slow potentials and the occipital EEG in congenital blindness. J Neurol Sci 37:51–58.

Olavarria JF, Hiroi R (2003) Retinal influences specify cortico-cortical maps by postnatal day six in rats and mice. J Comp Neurol (United States) 459:156–172.

Ossandón JP, Stange L, Gudi-Mindermann H, Rimmele JM, Sourav S, Bottari D, Kekunnaya R, Röder B (2023) The development of oscillatory and aperiodic resting state activity is linked to a sensitive period in humans. Neuroimage 275:120171.

Pachitariu M, Steinmetz N, Kadir S, Carandini M, Harris KD (2016) Kilosort: Realtime spike-sorting for extracellular electrophysiology with hundreds of channels. bioRxiv 061481.

Pelli DG (1997) The VideoToolbox software for visual psychophysics: Transforming numbers into movies. Spat Vis 10:437–442.

Poulet JFA, Crochet S (2019) The cortical states of wakefulness. Front Syst Neurosci 12:64.

Reimer J, Froudarakis E, Cadwell CR, Yatsenko D, Denfield GH, Tolias AS (2014) Pupil fluctuations track fast switching of cortical states during quiet wakefulness. Neuron (United States) 84:355–362.

Renart A, de la Rocha J, Bartho P, Hollender L, Parga N, Reyes A, Harris KD (2010) The asynchronous state in cortical circuits. Science (United States) 327:587–590.

Ribic A (2020) Stability in the face of change: Lifelong experience-dependent plasticity in the sensory cortex. Front Cell Neurosci 0:.

Riyahi P, Phillips MA, Colonnese MT (2021) Input-independent homeostasis of developing thalamocortical activity. eNeuro (United States) 8:10.1523/ENEURO.0184-Jun.

Röder B, Kekunnaya R (2022) Effects of early visual deprivation.

Rossant C, Kadir SN, Goodman DF, Schulman J, Hunter ML, Saleem AB, Grosmark A, Belluscio M, Denfield GH, Ecker AS, Tolias AS, Solomon S, Buzsaki G, Carandini M, Harris KD (2016) Spike sorting for large, dense electrode arrays. Nat Neurosci (United States) 19:634–641.

Saleem AB, Lien AD, Krumin M, Haider B, Rosón MR, Ayaz A, Reinhold K, Busse L, Carandini M, Harris KD (2017) Subcortical source and modulation of the narrowband gamma oscillation in mouse visual cortex. Neuron 93:315–322.

Seabrook TA, El-Danaf RN, Krahe TE, Fox MA, Guido W (2013) Retinal input regulates the timing of corticogeniculate innervation. J Neurosci (United States) 33:10085–10097.

Senzai Y, Fernandez-Ruiz A, Buzsáki G (2019) Layer-specific physiological features and interlaminar interactions in the primary visual cortex of the mouse. Neuron 101:500–513.e5.

Shen J, Colonnese MT (2016) Development of activity in the mouse visual cortex. J Neurosci (United States) 36:12259–12275.

Shinomoto S et al (2009) Relating neuronal firing patterns to functional differentiation of cerebral cortex. PLoS Comput Biol (United States) 5:e1000433.

Storchi R, Bedford RA, Martial FP, Allen AE, Wynne J, Montemurro MA, Petersen RS, Lucas RJ (2017) Modulation of fast narrowband oscillations in the mouse retina and dLGN according to background light intensity. Neuron 93:299–307.

Uhlhaas PJ, Haenschel C, Nikolić D, Singer W (2008) The role of oscillations and synchrony in cortical networks and their putative relevance for the pathophysiology of schizophrenia. Schizophr Bull 34:927–943.

Van Diepen RM, Foxe JJ, Mazaheri A (2019) The functional role of alpha-band activity in attentional processing: The current zeitgeist and future outlook. Curr Opin Psychol 29:229–238.

van Kerkoerle T, Self MW, Dagnino B, Gariel-Mathis M, Poort J, van der Togt C, Roelfsema PR (2014) Alpha and gamma oscillations characterize feedback and feedforward processing in monkey visual cortex. Proc Natl Acad Sci U S A 111:14332–14341.

Veit J, Hakim R, Jadi MP, Sejnowski TJ, Adesnik H (2017) Cortical gamma band synchronization through somatostatin interneurons. Nat Neurosci (United States) 20:951–959.

Veraart C, De Volder AG, Wanet-Defalque MC, Bol A, Michel C, Goffinet AM (1990) Glucose utilization in human visual cortex is abnormally elevated in blindness of early onset but decreased in blindness of late onset. Brain Res 510:115–121.

Vinck M, Batista-Brito R, Knoblich U, Cardin JA (2015) Arousal and locomotion make distinct contributions to cortical activity patterns and visual encoding. Neuron (United States) 86:740–754.

Wang B, Bernardez Sarria MS, An X, He M, Alam NM, Prusky GT, Crair MC, Huang ZJ (2021) Retinal and callosal activity-dependent chandelier cell elimination shapes binocularity in primary visual cortex. Neuron 109:502–515.e7.

Wen W, Turrigiano GG (2024) Keeping your brain in balance: Homeostatic regulation of network function. Annu Rev Neurosci.

Wu YK, Hengen KB, Turrigiano GG, Gjorgjieva J (2020) Homeostatic mechanisms regulate distinct aspects of cortical circuit dynamics. Proc Natl Acad Sci U S A (United States) 117:24514–24525.

